# Hypoxia induced lipid alterations in iPSC-derived human glial progenitor cells revealed by Matrix assisted laser desorption ionization mass spectrometry based cellular fingerprinting

**DOI:** 10.1101/2025.08.27.672639

**Authors:** Saira Hameed, Sivaramakrishnan Ramadurai, Maria E. Sanita, Daniel McGill, Felicia Green

## Abstract

**Background:** Increasing evidence suggests that the development of highly aggressive form of brain tumor “glioblastoma” contain cancer stem cells that closely resemble glial progenitor cells. The hypoxic microenvironment of tumors leads to metabolic reprogramming of cells, driving them towards more aggressive and treatment resistant state. To identify better targets for the effective removal of hypoxia adaptive cells, it is crucial to understand how these cells alter metabolism in hypoxic conditions.

**Methods:** We have used confocal microscopy to visualize mitochondrial morphology which confirmed hypoxia, and matrix assisted laser desorption ionization mass spectrometry imaging (MALDI-MSI) to decipher lipid metabolic changes in iPSC-derived human glial progenitor cells induced by that hypoxia.

**Results:** Our findings revealed that hypoxia induced changes in mitochondrial morphology, interaction of mitochondria with lysosomes, and the expression of oxidized lipid species in iPSC-derived human glial progenitor cells. Hypoxic cells showed elongated mitochondria that resembled strings. MALDI-MSI based finger printing showed upregulation of oxidized phosphatidylethanolamine (PE): *m/z* 716.484 PE(33:1)+O, *m/z* 730.01 PE(34:1)+O, *m/z* 766.497 PE(37:5)+O, *m/z* 768.510 PE(37:4)+O, and oxidized phosphatidylcholine (PC): *m/z* 742.500 PC(32:2)+O, *m/z* 744.513 PC(33:2)+O, *m/z* 770.531 PC(34:2)+O), *m/z* 794.534 PC(34:1)+O, *m/z* 888.523 PC(42:11)+OH, and oxidized phosphatidic acid (PA): *m/z* 673.444 PA(33:2)+O, *m/z* 699.466 PA(35:3)+OH, *m/z* 725.531 PA(37:4)+OH in hypoxic cells.

**Conclusions:** Given the importance of glial progenitor cells in the central nervous system, as a precursor of glioblastoma, and their structural reliance on lipids, the molecular perturbations in the cell as a result of oxygen deficiency (hypoxia) remains unclear. The change in mitochondrial morphology showed that the cells were under hypoxic stress and mass spectrometry imaging unveiled important lipid metabolic changes in hypoxic iPSC-derived human glial progenitor cells as byproducts of oxidative stress, and provided insights that could lead to better treatment strategies for hypoxia-resistant cells.

Graphical Abstract

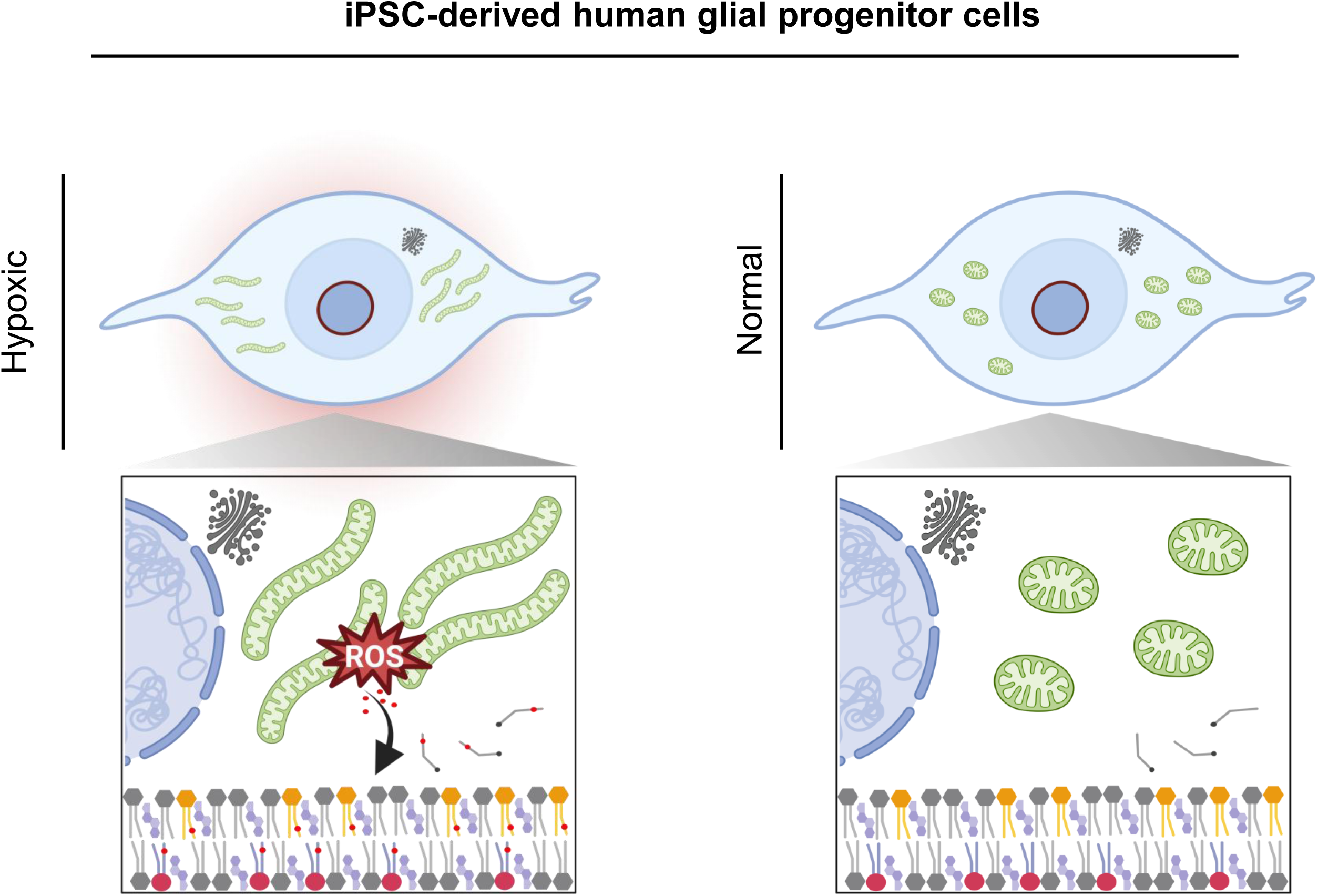

**Graphical abstract** was created in BioRender https://app.biorender.com/illustrations/6822efb6aadd8c37669dd403

## Background

According to the World Health Organization (WHO) glioblastoma is the most common and aggressive form of brain tumor [1]. Growing body of evidence suggests that glial progenitor cells are precursors of glioblastoma [2]. Oxygen deficiency (hypoxia) is observed in glioblastoma lesions [3], that causes disruption of blood brain barrier [4], promotes drug resistance, invasion and inhibition of immune response [5]. Hypoxia impacts mitochondrial activity, lipid metabolism, and contributes to cellular stress and cytotoxicity [6]. Interestingly, hypoxia is associated with activation of glial cells that release cytokines and reactive oxygen species [7]. Pharmaceutical interventions or genetic approaches to block glial cell activation relieved ischemia that confirmed the critical role of these cells in the expression of damage during oxygen deficiency in the brain [8]. Despite the studies mentioned above, hypoxia induced lipidomic profiling of iPSC-derived human glial progenitor cells at single cell level remains elusive.

Previous studies have demonstrated the impact of hypoxia on lipid accumulation in mouse fibroblast cells, and cervical cancer derived (HeLa) cells by biochemical assays of total lipid extracts [9, 10], in pancreatic cancer cells by Ramen scattering microscopy [11]. The conventional methods suffer some limitations therefore in this study we used Matrix assisted laser desorption ionization mass spectrometry imaging (MALDI-MSI) that allows label free detection, mapping, and quantification of a variety of biomolecules [12]. Recent studies have shown lipid imaging with MALDI-MSI [13, 14] in brain pathologies [15], and cultured cells [16].

In this study, mitochondrial morphology was evaluated by confocal microscopy and lipid alterations by single cell MALDI-MSI fingerprinting. The model provided a valuable tool for detection and characterization of lipids associated with hypoxic stress in iPSC-derived human glial progenitor cells, which in pathological conditions such as glioblastoma can serve as biomarkers.

## Results

### Glial progenitor cells under stress

#### Effects of hypoxia on mitochondria determined by confocal microscopy

Mitochondrial structure is controlled by cell activities [17]. Studies have shown that mitochondrial respiration and morphology are impaired by hypoxia [18]. Here we show using confocal microscopic evaluations that the mitochondria appear elongated resembling strings in the hypoxic cells (Fig 1A) in agreement with previous studies [19], whereas mitochondria appear, as expected, oval in normal cells (Fig 1B).

**Fig. 1.**
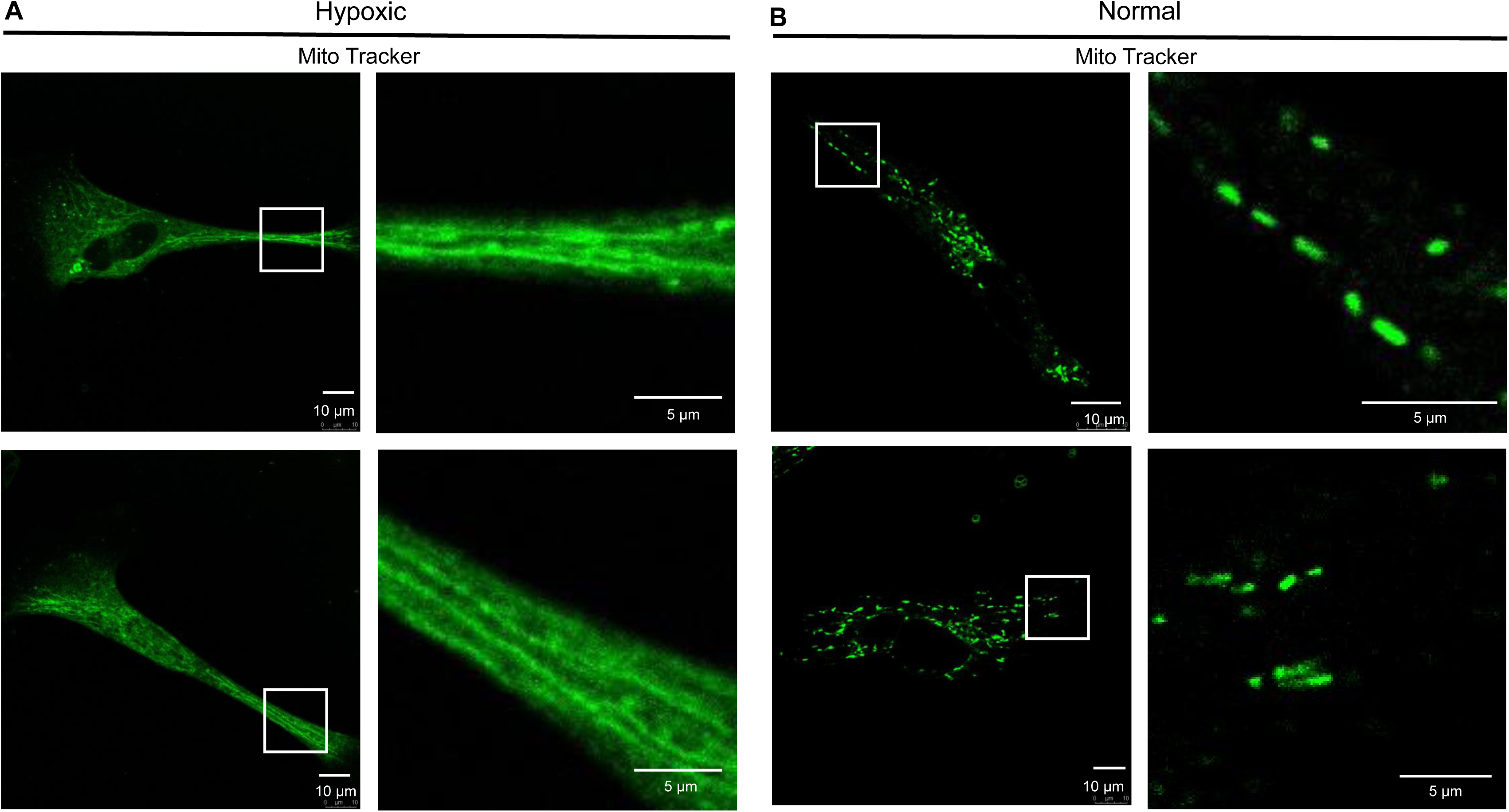
Hypoxia reshapes mitochondria in glial progenitor cells. The cells were stained with Mito Tracker green dye and were observed by confocal microscopy. **A** Hypoxic cells showed elongated mitochondria resembling strings. **B** Normal cells showed oval shaped mitochondria. Scale bars 5 µm, and 10 µm.

### Mitochondrial and lysosomal interaction in glial progenitor cells under hypoxic stress

Mitochondria and lysosome interact through physical and functional connections to regulate metabolism and manage cellular stress during hypoxia in MKN45 cells (gastric cancer cell line) [20], and Hela cells (cervical cancer cell line) [21]. In this study we tested the hypothesis and compared the hypoxic and normal iPSC-derived human glial progenitor cells. Both were double stained with Mito Tracker (green fluorescent mitochondria marker) and Lyso Tracker (red fluorescent lysosome marker) to allow us access to the interaction between mitochondria and lysosome. Hypoxic cells showed more prominent lysosomes (red) and their overlay with mitochondria (green) produced yellow signal in regions of signal overlap (Fig. 2A) whereas, although the normal cells had lysosome signals (red) in the same regions as mitochondria (green) they were weaker and did not show up yellow in the overlay. (Fig. 2B).

**Fig. 2.**
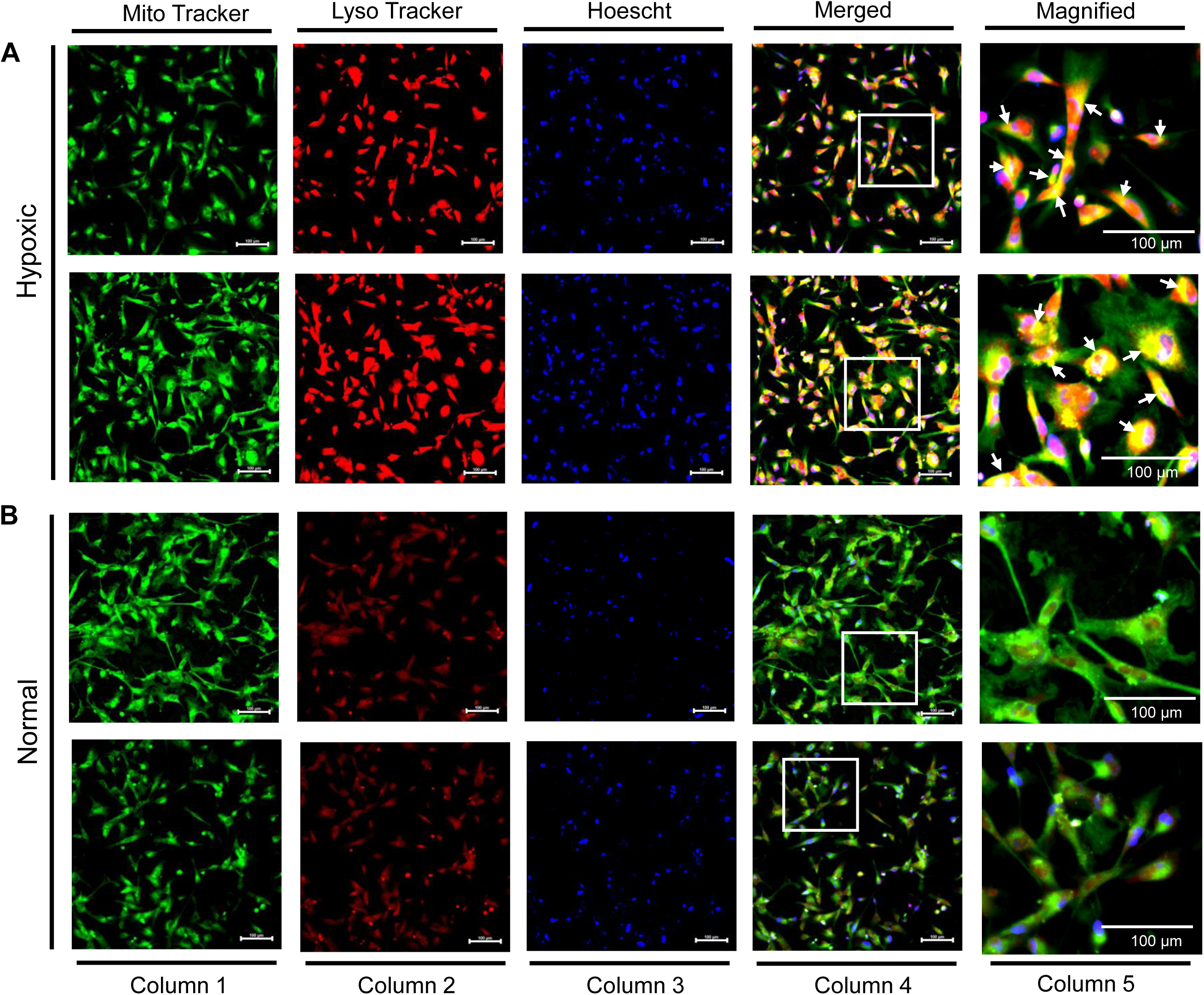
Hypoxia promotes interaction between mitochondria and lysosomes in glial progenitor cells. The cells were stained with Mito Tracker green (mitochondrial stain), Lyso Tracker red (lysosome stain), Hoechst blue (nucleus stain), and were observed by fluorescence microscopy. Column 1 shows mitochondria, column 2 shows lysosomes, column 3 shows nuclei, column 4 shows the overlay, and column 5 shows the expanded region outlined in column 4. Figures in **A** are two examples of hypoxic cells that showed yellow signal in the regions of interactions between mitochondria and lysosomes, as compared to figures in **B** that shows two examples of normal cells with no yellow signal. Mitochondria (green), lysosome (red), nucleus (blue). Scale bar 100 µm at 20x magnification.

### Single cell imaging mass spectrometry

SEM results showed that small matrix crystals are completely volatilized by laser shot during MALDI-MSI (Supplementary Fig. 1A and 1B) [22]. Moreover, small matrix crystals allowed us access to higher spatial resolution MALDI MSI of ∼ 5 µm (Supplementary Fig. 1C and 1D) [23]. This enabled us to clearly define cell boundaries within the MALDI MSI and obtain MS from an individual cell, as shown in Figure 3.

**Fig. 3.**
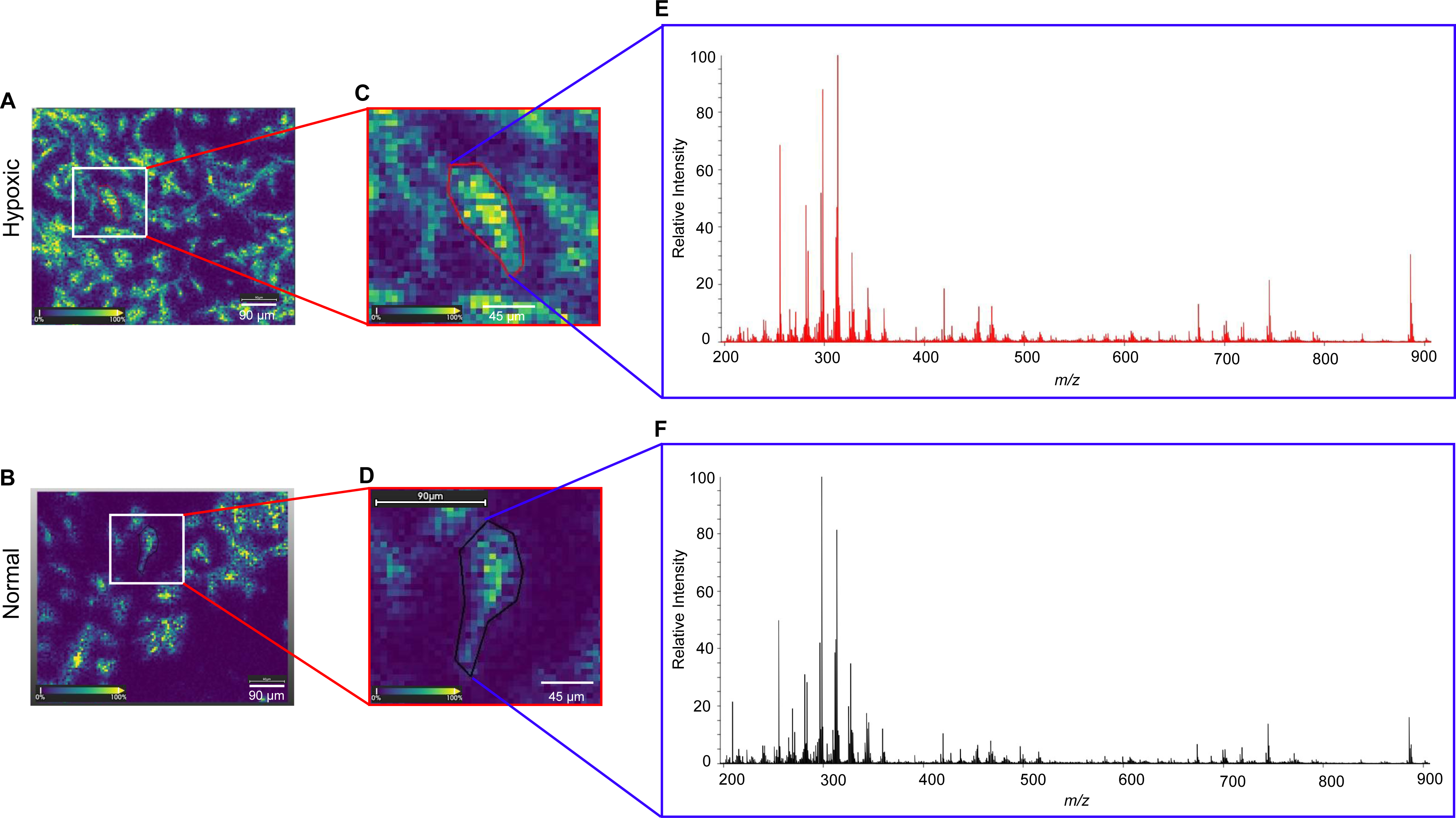
Hypoxia promotes molecular alterations in glial progenitor cells. The cells were analyzed by MALDI-MSI for molecular fingerprinting. Figure **A** shows ion image of hypoxic cells. **B** shows normal cells. Regions of interest were drawn around single cells from hypoxic and normal ion images. **C** shows enlarged hypoxic cell encircled with red line, and **D** shows normal cell encircled with black line. Single cell mass spectra were extracted from the selected regions of interest. Fig **E** shows single cell mass spectrum (red) from the hypoxic cell, and **F** shows single cell mass spectrum (black) from the normal cell. Scale bar in A and B are 90 µm, and C and D are 45 µm.

Figure 3A and 3B shows ion images of hypoxic and normal iPSC-derived human glial progenitor cells. Regions of interest (ROIs) were drawn around individual hypoxic (red line) and normal (black line) cells. Fig. 3C and 3D shows enlarged cells with ROIs. Fig 3E shows red mass spectrum from hypoxic cells, and Fig 3F shows black spectrum from normal cell. By repeating this process we were able to obtain an array of MS from all the cells analysed, both 180 hypoxic and 180 normal. This allows us to molecularly profile hypoxic and normal iPSC-derived human glial progenitor cells. During hypoxic stress, cells undergo lipid remodelling to adapt to low oxygen levels [6]. MALDI-MSI showed hypoxia induced changes in the expression of lipids in glial progenitor cells (Supplementary Table 1). The calculations of upregulation and downregulation in 48 hours hypoxic and normal glial progenitor cells were based on *p* values and percentage differences in peak intensities between hypoxic and normal cells (as shown in supplementary table 1). The *p* values for all the mass peaks were below 0.001 suggesting that changes in intensity between the hypoxic and normal cells were statistically significant, although it is also somewhat indicative of the large (180 cells in each group) data set used. Interestingly, MALDI-MSI showed up regulation of over 50% for 29 peaks and downregulation of over 50% for 38 peaks out of the top 76 most intense peaks in the mass spectra. Out of these peaks we were able to confirm identification of 12 oxidized lipids through the use of MALDI - MS/MS directly from the cells (supplementary Figure 2). These were oxidized phosphatidylethanolamine (PE) (Fig. 4): *m/z* 716.484 PE(33:1)+O, *m/z* 730.01 PE(34:1)+O, *m/z* 766.497 PE(37:5)+O, *m/z* 768.510 PE(37:4)+O, and oxidized phosphatidylcholine (PC) (Fig. 5): *m/z* 742.500 PC(32:2)+O, *m/z* 744.513 PC(33:2)+O, *m/z* 770.531 PC(34:2)+O), *m/z* 794.534 PC(34:1)+O, *m/z* 888.523 PC(42:11)+OH, and oxidized phosphatidic acid (PA) (Fig. 6): *m/z* 673.444 PA(33:2)+O, *m/z* 699.466 PA(35:3)+OH, *m/z* 725.531 PA(37:4)+OH and all showed upregulation in hypoxic cells as compared to normal cells. Molecules that weren’t firmly identified but showed >50% change in peak intensity and a *p*-value of less than 0.001 are shown in Supplementary Figure 3 for molecules downregulated by hypoxia and in Supplementary Figure 4 for molecules upregulated by hypoxia. Some of these have been given putative assignments through database searches and these are shown in Supplementary Table 1.

**Fig. 4.**
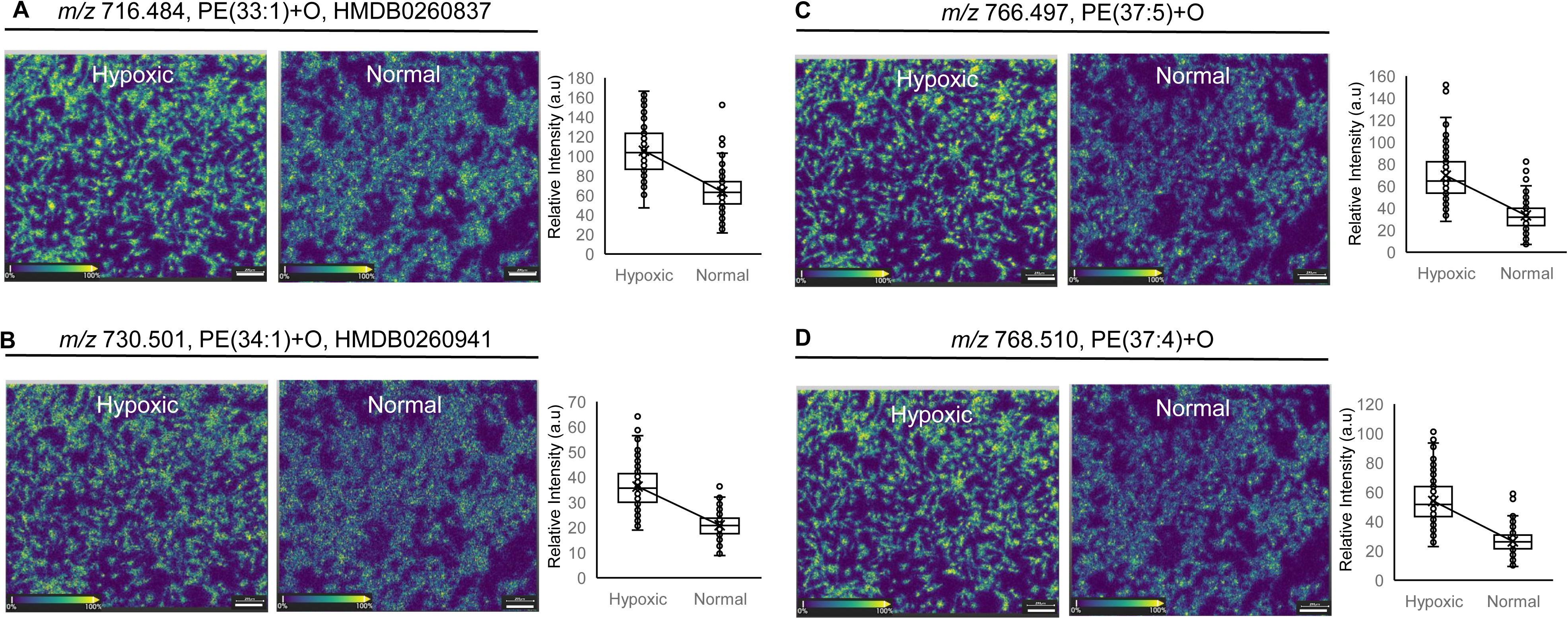
Hypoxia promotes oxidation of phospholipids in glial progenitor cells. MALDI-MSI of the cells showed upregulation of oxidized phosphatidylethanolamines (OxPE) in hypoxic cells. Example images from MALDI-MSI of hypoxic and normal cells and whisker plots containing all data at **A** *m/z* 716.484 PE(33:1)+O, HMDB0260837. **B** *m/z* 730.01 PE(34:1)+O, HMDB0260941. **C** *m/z* 766.497 PE(37:5)+O. **D** *m/z* 768.510 PE(37:4)+O. Scale bar 200 µm. Box and Whisker plots were drawn in Excel with the data collected from 6 measurement regions and 180 cells (hypoxic and normal) each.

**Fig. 5.**
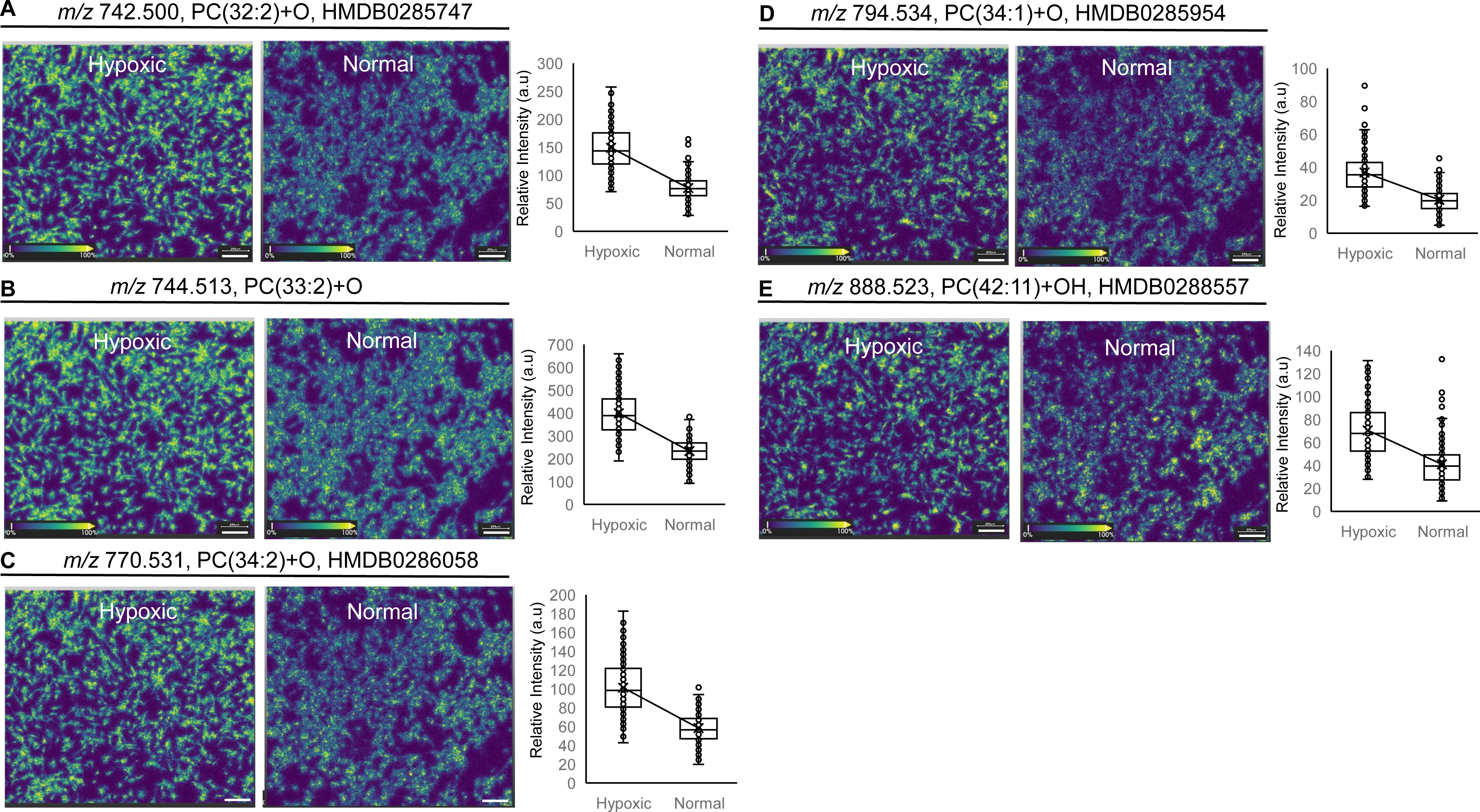
Glial progenitor cells showed changes in metabolism caused by hypoxia. Example images from MALDI-MSI of hypoxic and normal cells and whisker plots containing all data MALDI-MSI of the cells showed upregulation of Oxidized phosphatidylcholine (OxPC) in hypoxic cells. **A** *m/z* 742.500 PC(32:2)+O, HMDB0285747. **B** *m/z* 744.513 PC(33:2)+O. **C** *m/z* 770.531 PC(34:2)+O, HMDB0286058. **D** *m/z* 794.534 PC(34:1)+O, HMDB0285954. **E** *m/z* 888.523 PC(42:11)+OH, HMDB0288557. Scale bar 200 µm. Box and Whisker plots were drawn in Excel with the data collected from 6 measurement regions and 180 cells (hypoxic and normal) each.

**Fig. 6.**
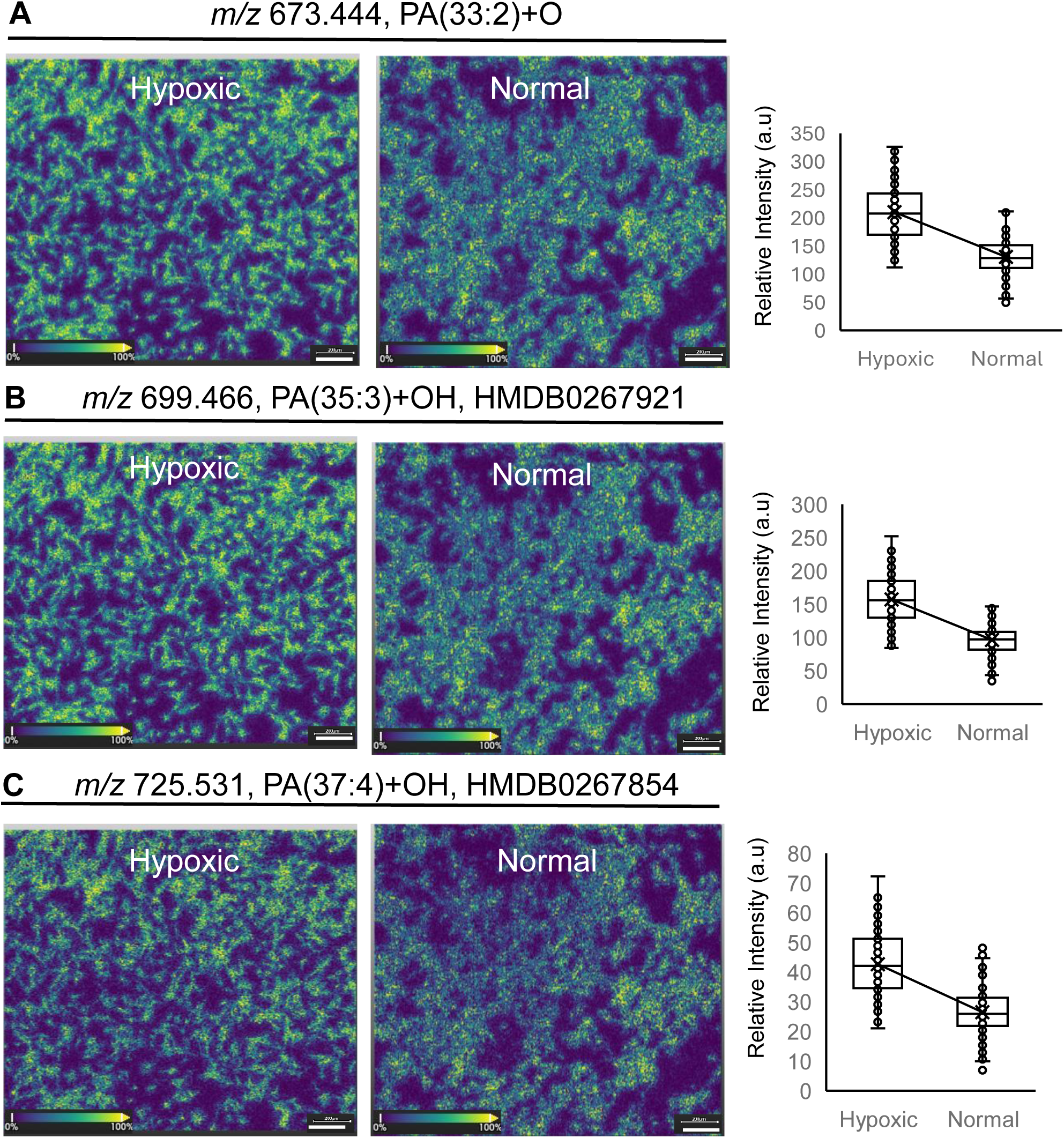
Hypoxia causes oxidation of fatty acids. MALDI-MSI of glial progenitor cells showed Oxidized phosphatidic acid (OxPA) upregulation in hypoxic cells. Ion images of hypoxic and normal cells and whisker plots at **A** *m/z* 673.444 PA(33:2)+O. **B** *m/z* 699.466 PA(35:3)+OH, HMDB0267921. **C** *m/z* 725.531 PA(37:4)+OH, HMDB0267854. Scale bar 200 µm. Box and Whisker plots were drawn in Excel with the data collected from 6 measurement regions and 180 cells (hypoxic and normal) each.

## Discussion

Glia are immune cells of the nervous system that are triggered by stress stimuli and undergo remarkable phenotypic, molecular and operational changes [24]. In this study, we present a MALDI-MSI based cellular fingerprinting approach to explore hypoxia associated lipid markers in iPSC-derived glial progenitor cells. To the best of our knowledge, there is no previous report on the use of MALDI-MSI to interrogate lipid alterations in stressed glial progenitor cells at single cell level. We choose to study glial progenitors under hypoxic stress because of the association of such cells and the hypoxic environment with glioblastoma that is the most aggressive form of brain tumour [2].

Together, the current findings explore the molecular differences of cells that have reached a hypoxic state as revealed by the elongated mitochondria in hypoxic cells (Fig 1A) as compared to oval shaped mitochondria seen in normal cells. (Fig 1B). Based on earlier studies the formation of long strings of interconnected mitochondria in the shape of tubular nanotunnels (referred as “mitochondria-on-a-string”) are induced in hypoxic mouse brains through inter-mitochondrial contact and fusion [25]. Therefore, the change in basic morphology of the mitochondrial organelles from that in normal cells (generally considered as oval-like structure) confirms hypoxia [26].

Exposure of cells to hypoxia causes oxidative stress [27], and generation of reactive oxygen species (ROS) as toxic byproducts [28]. The mitochondrial electron transport chain undergoes series of reactions in which molecular oxygen is reduced to superoxide radical (O_2_ ^•−^), and hydrogen peroxide (H O) by superoxide-dismutase [29]. Superoxide and hydrogen peroxide are the primary reactive oxygen species (ROS) that undergo series of chain reactions to form more reactive secondary species including ^•^OH radicals [29]. The ROS damage building blocks and lead to cellular dysfunction. Therefore, a mitochondrial quality control system is activated during hypoxia to suppress the accumulation of ROS. Lipids appear as primary barriers to the free circulation of ROS between cells and become targets of oxidation [30]. The oxidation reactions initiated by ROS can produce lipid radicals that can propagate formation of other oxidized lipids through reactions involving isomerisation and chain cleavage [30]. Oxidized lipids accumulate in apoptotic cells and microparticles released by activated and dying cells [31], and are considered as lipotoxic factors that are ubiquitous and found in several inflammatory settings including neuroinflammation [32].

Lipids have long been recognized to accumulate in cells under hypoxic stress [9], and therefore were the target of the study. Our single cell MALDI-MSI approach was capable of detecting hypoxia induced upregulation of on average 88% ± 5% for oxidized phosphatidylethanolamine (specifically PE: *m/z* 716.484 PE(33:1)+O, *m/z* 730.01 PE(34:1)+O, *m/z* 766.497 PE(37:5)+O, *m/z* 768.510 PE(37:4)+O – see Figure 4), in glial progenitor cells. This could be caused by the hypoxia pushing the cell towards cell death as a recent study showed that the expression of oxidized PE could be linked as a key “eat-me” signal on ferroptotic cells surface [33]. However, hypoxia can regulate ferroptosis in specific cells and conditions through different pathways [34], that will be explored in future studies.

MALDI-MSI of hypoxic glial progenitor cells also showed upregulation of on average 78% ± 9% for oxidized phosphatidylcholine (PC) (specifically *m/z* 742.500 PC(32:2)+O, *m/z* 744.513 PC(33:2)+O, *m/z* 770.531 PC(34:2)+O, *m/z* 794.534 PC(34:1)+O, *m/z* 888.523 PC(42:11)+OH as shown in Figure 5). Previous studies have shown a relationship between oxidized PC and the pathogenesis of various oxidative stress related diseases of the nervous system [32]. This study confirms the expected upregulation of oxidized PCs in glial progenitor cells when under oxidative stress caused by hypoxia.

Moreover, glial progenitor cells under hypoxic stress also showed upregulation of on average 62 ± 2% for oxidized phosphatidic acids (PA) (specifically *m/z* 673.444 PA(33:2)+O, *m/z* 699.466 PA(35:3)+OH, *m/z* 725.531 PA(37:4)+OH as shown in Figure 6). In this study upregulation of oxidized PAs is observed for the first time in hypoxic glial progenitor cells, and may be considered as a novel bioactive molecule [35].

Surprisingly, the oxidized PE, oxidized PC and oxidized PA detected in glial progenitor cells induced by hypoxic stress were lipid species which have not been specifically reported by previous studies. This may be because the commonly used methods to monitor hypoxia associated lipids were not MS but rather light microscopy [9], biochemical assay based on total lipid extraction [10], Raman spectroscopy [11], and immunoassays based on antibody staining [36]. These conventional methods suffer some limitations, light microscopy struggles to provide comprehensive information about cellular lipids due to limited resolution and inability to monitor lipid structures [37]. The total lipids extraction methods disrupt the native cellular environment and therefore can underestimate certain molecules due to difficulty in extraction [38]. Raman spectroscopy suffers from weak signal intensities so may not be comprehensive [39]. While, antibody staining for cellular lipid analysis suffer because lipids are typically poor immunogens [40]. Although some antibodies can detect head groups of certain oxidized lipids but cannot distinguish them based on their structures [36]. Whereas our MALDI-MSI based cellular finger printing strategy is an excellent method to detect lipids. We have shown here it’s potential as a tool to investigate hypoxic stress in individual glial progenitor cells and detect hypoxia associated metabolic perturbations for single cells. A key feature of our untargeted approach is that it enables rapid detection of novel oxidized lipid species induced by hypoxic stress in glial progenitor cells. This allows us to, in combination with tandem mass spectrometry, to find and identify markers and byproducts of oxidative stress in an untargeted and single cell manner. This could provide molecular insights that could lead to better treatment strategies for hypoxia-resistant cells.

**Supplementary Fig. 1 Secondary electron microscopy (SEM) of spray coated 1,5-diaminonaphthalene (DAN) matrix, and raster scanned region. A** SEM image of DAN crystals. Scale bar 1.64 µm, 1.58 µm, 2.071 µm. **B** SEM image of raster scanned region with laser irradiation by MALDI-MSI. Scale bar 4.549 µm, 4.186 µm. **C** MALDI-MSI images of glial progenitor cells at *m/z* 324.745, and *m/z* 496.601. Scale bar 90 µm, 45 µm, 22.5 µm, 11.25 µm. Selected pixels encircled and pointed with white arrow ∼ 5 µm.

**Supplementary Fig. 2** Tandem mass spectra of identified molecules at **A** *m/z* 766. 497, PE(20:4/17:1-O). **B** *m/z* 730.501, PE(34:1)+O, HMDB0260941, **C** *m/z* 766.497, PE(37:5)+O, **D** *m/z* 768.510, PE(37:4)+O, **E** *m/z* 742.500, PC(32:2)+O, HMDB0285747, **F** *m/z* 744.513, PC(33:2)+O, **G** *m/z* 770.531, PC(34:2)+O, HMDB0286058, **H** *m/z* 794.534, PC(34:1)+O, HMDB0285954, **I** *m/z* 888.523, PC(42:11)+OH, HMDB0288557, **J** *m/z* 673.444, PA(33:2)+O, **K** *m/z* 699.466, PA(35:3)+OH, HMDB0267921, **L** *m/z* 725.531, PA(37:4)+OH, HMDB0267854. Putative identification from the resulting tandem mass (MS/MS) spectra was conducted in ChemDraw with reference to literature [41]. Tandem mass spectra of unidentified molecules at **M** *m/z* 274.950, **N** *m/z* 279.794, **O** *m/z* 310.783, **P** *m/z* 436.645, **Q** *m/z* 462.820, **R** *m/z* 645.416, **S** *m/z* 647.430, **T** *m/z* 772.541, **U** *m/z* 776.513, **V** *m/z* 792.519, **W** *m/z* 885.597, **X** *m/z* 886.504, **Y** *m/z* 887.513, **Z** *m/z* 911.514.

**Supplementary Figure 3 Alterations of unidentified molecules caused by hypoxia in glial progenitor cells.** Ion images of hypoxic and normal cells and whisker plots of the molecules downregulated by hypoxia at **A** *m/z* 274.950 (unidentified), B *m/z* 279.794 (unidentified), C *m/z* 310.783 (unidentified), **D** *m/z* 436.645 (unidentified), **E** *m/z* 462.829 (unidentified). Box and Whisker plots were drawn in Excel with the data collected from 180 cells (hypoxic and normal) each.

**Supplementary Figure 4 Alterations of unidentified molecules caused by hypoxia in glial progenitor cells.** Ion images of hypoxic and normal cells and whisker plots of the molecules upregulated by hypoxia at **A** *m/z* 645.427 (unidentified), **B** *m/z* 647.430, **C** *m/z* 772.541, **D** *m/z* 776.513, **E** *m/z* 792. 519, **F** *m/z* 885.597, **G** *m/z* 886.504, **H** *m/z* 887.513, **I** *m/z* 911.514. Scale bar 200 µm. Box and Whisker plots were drawn in Excel with the data collected from 180 cells (hypoxic and normal) each.

**Supplementary Table 1** Comparison of molecular profiles between normal and hypoxic glial progenitor cells.

## Materials and methods

### Chemicals

Reagents (analytical grade) used in the study were from Merck (Sigma-Aldrich, Darmstadt, Germany), and Fisher Scientific, United Kingdom, except where specifically noted. Good laboratory practices (GLP) were used to store, handle, and dispose of the chemicals used in the study.

### Maintenance of iPSC-derived human glial progenitor cells

iPSCs-derived human glial progenitor cells (derived from human skin fibroblasts) were generated by Dr. Selina Wray’s lab at University College London (UCL), with approval by the National Hospital for Neurology and Neurosurgery and Institute of Neurology Joint Research Ethics Committee: 09/H0716/64 [42], and kindly provided to us for research. The cells were received from a single clone at passage 9 (139 days) *in vitro* and were re-cultured until passage 16 (170 days) *in vitro* to use in the experiments. The cells were maintained in specialized cell culture N2B27 media as outlined in the previous study [42]. Briefly, one hundred millilitre N2B27 media containing 48.4ml Dulbecco’s modified eagle medium/Ham’s F-12 (DMEM/F12) + Glutamax (1X) (Gibco), 48.5ml Neurobasal Media (Gibco), 500uL N-2 Supplement (Gibco), 1ml B-27 Supplement (Gibco), 500uL L-Glutamine, 500uL Non-essential amino acids (Gibco), 100uL B-mercaptoethanol, 500uL PenStrep, 25uL Insulin, with an addition of fibroblast growth factor-2 (FGF2) at 10ng/ml (Peprotech). Prior to seeding the cells, culture plates were coated with Geltrex (Gibco) and incubated for 1 hour at 37°C, with 5% CO_2_. The cells were grown in the humidified incubator at 37°C, with 5% CO_2_, with medium changed twice weekly, until ready to passage at ∼80-90% confluency. Detachment of the cells was carried out with incubation of the cells in 0.5mM ethylenediaminetetraacetic acid (EDTA) diluted in Dubecco’s phosphate buffered saline (DPBS) buffer (without Ca^2+^ and Mg^2+^ (Gibco); the same DPBS was used for washing prior to this step. Upon detachment, the EDTA was removed and the cells were washed from the bottom of the plate with fresh N2B27 media, to be re-plated at a 1:3 ratio.

### Cell culture and hypoxic treatments

Cells suspended in culture medium were cultured directly onto polished soda lime, float glass slides, with dimension 25mm X 75mm, thickness 1.1 mm, pre-coated with fully oxidized indium-tin-oxide, ITO thickness 100 nm, ITO resistance 16-20 Ω/sq (Ossila, UK). Silicon well separators were used to make temporary chambers onto the glass surface. Approximately, 0.1 x 10^6^ cells in 0.5 ml of culture medium were directly seeded into each chamber of the slide and grown for optimal confluence [14]. Cells were subjected to hypoxic conditions for 48 h using nBIONIX-1 hypoxic cell culture kit (Xcelitis, GmbH, Mannheim, Germany) as reported before [43]. Briefly, an oxygen meter, petri dish with cell culture slide, and gas concentration (oxygen absorber) agent were placed in a gas barrier pouch that was sealed. The gas concentration agent was isolated with a clip when the oxygen concentration reached the desired levels. The oxygen levels were constantly monitored with an oxygen meter inside the kit and remained stable throughout experiments. Hypoxic cells were grown in 4% oxygen to maintain self-renewal and limit spontaneous differentiation [44], and normal cells were grown in 20.9% oxygen. In this study hypoxia was induced in glial progenitor cells without any chemical treatment. Because introduction of chemical inducers can effect cellular functions apart from inducing hypoxia [45].

### Washing and drying

The cell culture slides were observed under the light microscope and were washed with phosphate buffer saline (PBS) (Fisher scientific) buffer, followed by deionized water and 150mM solution of ammonium acetate (Fisher scientific). After washing the slides were dried in a vacuum desiccator.

### Paraformaldehyde fixation

The slides were chemically fixed in 4% paraformaldehyde (PFA) solution prepared in PBS buffer for around 10 min at room temperature. After fixation the slides were washed following the protocol described above.

### Confocal microscopy

Glial progenitor cells were seeded in 8 well chambered cover slides (Ibidi GmBH) (∼25000 cells per well). Cells were grown under hypoxic and normal conditions as described above. The hypoxic and normal cell slides were chemically fixed in 4% PFA solution prepared in PBS for around 10 min at room temperature. After fixation the slides were washed and labelled with Mito Tracker green (stained mitochondria), Lyso Tracker red (stained lysosome) and Hoescht (stained nucleus) for 10 minutes. The slides were washed and imaged by using Leica DMi8 laser scanning confocal microscope at 63x magnification (oil immersion objective) with laser excitation 405, 488, 568nm for Hoescht, Mito Tracker green and Lyso Tracker red respectively.

### Optical microscopy

After formalin fixation the slides were labelled with Mito Tracker green (stained mitochondria), Lyso Tracker red (stained lysosome) and Hoescht (stained nucleus) for 10 minutes. The slides were washed, dried in a vacuum desiccator, and observed under the fluorescent microscope (Leica, Maheim, Germany) using 20x objective.

### Matrix coating

The slides were spray coated with 1,5-diaminonaphthalene 97% (DAN) (Sigma-Aldrich) 5mg/ml dissolved in 90% acetonitrile using HTX M3+ sprayer (HTX Technologies LLC, USA). Temperature of the spray nozzle was set at 80 °C, pressure at 10 psi, flow rate 100 µL/min, velocity 1200 mm/min, track spacing 2 mm, number of passes 8, and drying time 10 sec.

### Matrix assisted laser desorption ionization mass spectrometry imaging (MALDI-MSI)

Normal and hypoxic glial progenitor cells cultured on different slides were analysed simultaneously using the two-chamber slide holder (Bruker Daltonics). MALDI MSI data was acquired on a tims-TOF fleX system (Bruker Daltonics) equipped with a smart-beam 3D 10□kHz Nd:YAG laser (355 nm) and micro-GRID, controlled by TimsControl v4.1 and flexImaging v7.2 software (Bruker Daltonics). Data was acquired in negative ion mode, *m/z* range of 150–2000, with 20 laser shots per pixel, 10□kHz laser frequency, laser spot size 5□µm^2^ and raster 5□µm as reported before [46, 47]. The tune parameters were the following: MALDI Plate Offset 50□V, Deflection 1 Delta −70 V, Funnel 1 RF 300 Vpp, isCID Energy 0□eV, Funnel 2 RF 400 Vpp, Multipole RF 500 Vpp, Collision Energy 10□eV, Collision RF 800 Vpp, Quadrupole Ion Energy 10□eV and Low Mass 20□m/z, Focus Pre TOF Transfer Time 70□µs and Pre Pulse Storage 7□µs, and Detection set to Focus Mode.

### Scanning electron microscopy

The crystal size of the spray coated matrix ≤ 2 µm (Supplementary Fig. 1A), and the rastered scanned region (Supplementary Fig. 1B) were measured by scanning electron microscopy (SEM).SEM photomicrographs were collected on a JSM-IT100 InTouchScope Scanning Electron Microscope (JEOL) with a secondary electron detector (SED) set at an accelerating voltage of 5 kV, a working distance of 12 mm, and a probe current of 40 (arbitrary units) under high vacuum. The spatial resolution of MALDI-MSI ion images were estimated from the pixel size ∼5 µm (Supplementary Fig. 1C and 1D).

### Data analysis and molecular assignments

Data was imported into SCiLS Lab MVS, Version 2025b Pro, from Bruker Daltonics, using the standard import parameters. Regions of interest (ROI) were drawn around 180 individual cells to extract mass spectra from six measurement regions of hypoxic and normal samples each. Each replicate was just re-expansion without any modification. Data was normalized to root mean square (RMS). This study was targeted on lipidomic fingerprinting of glial progenitor cells. Single cell mass spectra were scanned peak by peak manually at interval width ±10 ppm, between the range *m/z* 250 to *m/z* 1000, and ion images were saved. The feature table was exported to CSV format, converted to Excel for calculations. The data was expressed as mean ± standard deviation (SD) across 180 cells for both hypoxic and normal cells. The difference between hypoxic and normal groups were analysed using unpaired t-test with 2 tails and graphs were plotted using Excel functions. The mass peaks that showed differences in the ion images and from the calculated *p*-values were subjected to tandem mass spectrometry for identification and were shown in the paper. Tandem mass spectrometry was performed with random mode, partial sample, 50 shots at raster position, diameter limit 2000 µm, isolation width 1 *m/z*, and 3 *m/z*, collision energy 5 eV, 20 eV, 30 eV, 40 eV, and 50 eV. Putative identification from the resulting tandem mass (MS/MS) spectra was conducted in ChemDraw with reference to literature [41], HMDB database, and CFM-ID [48].

## Contributions

SH and SR conceived the study. MES and SR cultured cells and induced hypoxia. SR and SH collected confocal microscopy data. SH collected MALDI-MSI data. SH performed tandem mass spectrometry and DM performed structural elucidation of lipids with literature support. FG was supervisor. SH interpreted the results and wrote the manuscript with input from all the coauthors.

## Acknowledgements

Dr. Anna Warren and Dr. Trincao Jose performed scanning electron microscopy at the Diamond Light Source, Harwell Science and Innovation Campus, Didcot, Oxfordshire OX11 0DE, UK.

Dr. Anthony Devlin, Dr. Mackay Logan, Dr. Siyu Liu, and Dr. Kenny Robinson provided general lab support.

The Rosalind Franklin Institute is funded by UK Research and Innovation through the Engineering and Physical Sciences Research Council (ESPRC).

## Corresponding author

Correspondence to Dr. Saira Hameed, and Dr. Felicia Green

