## Supplementary figures and images for "Hypoxia induced lipid alterations in iPSC-derived human glial progenitor cells revealed by Matrix assisted laser desorption ionization mass spectrometry based cellular fingerprinting"

### Supplementary Figure 1

#### Supplementary Fig. 1

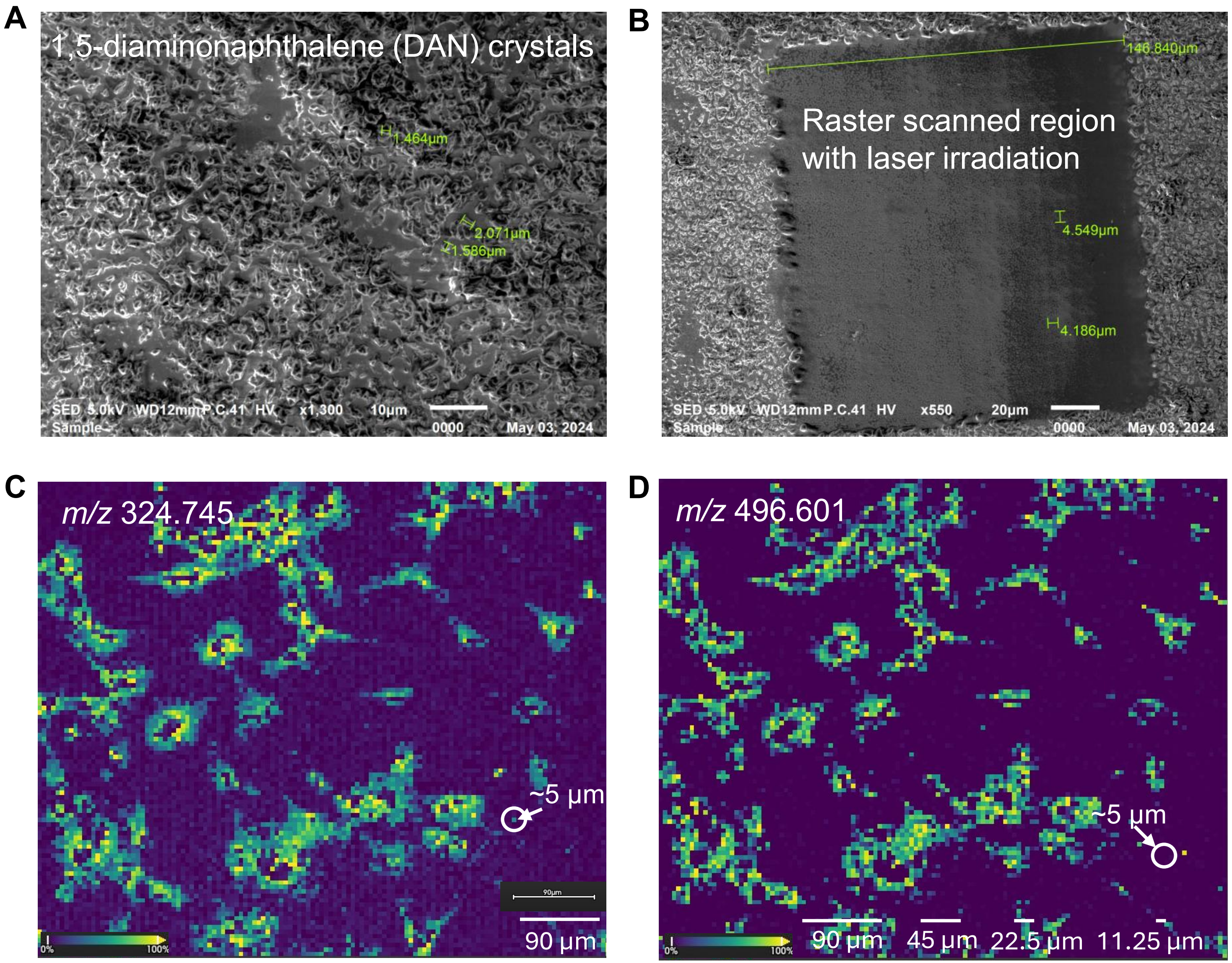

### Supplementary Figure 3

**Supplementary Fig. 3**

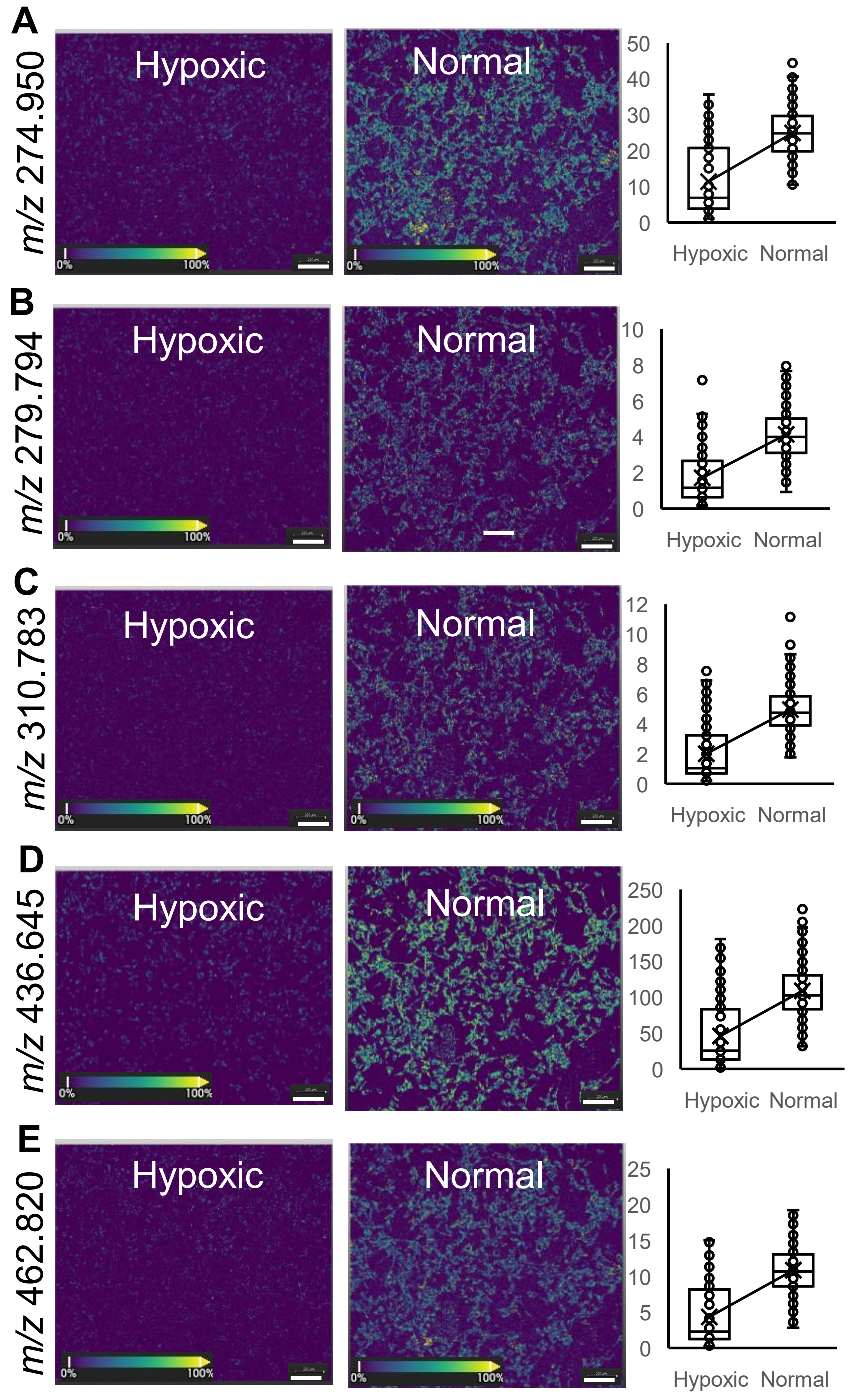

### Supplementary Figure 4

**Supplementary Fig. 4**

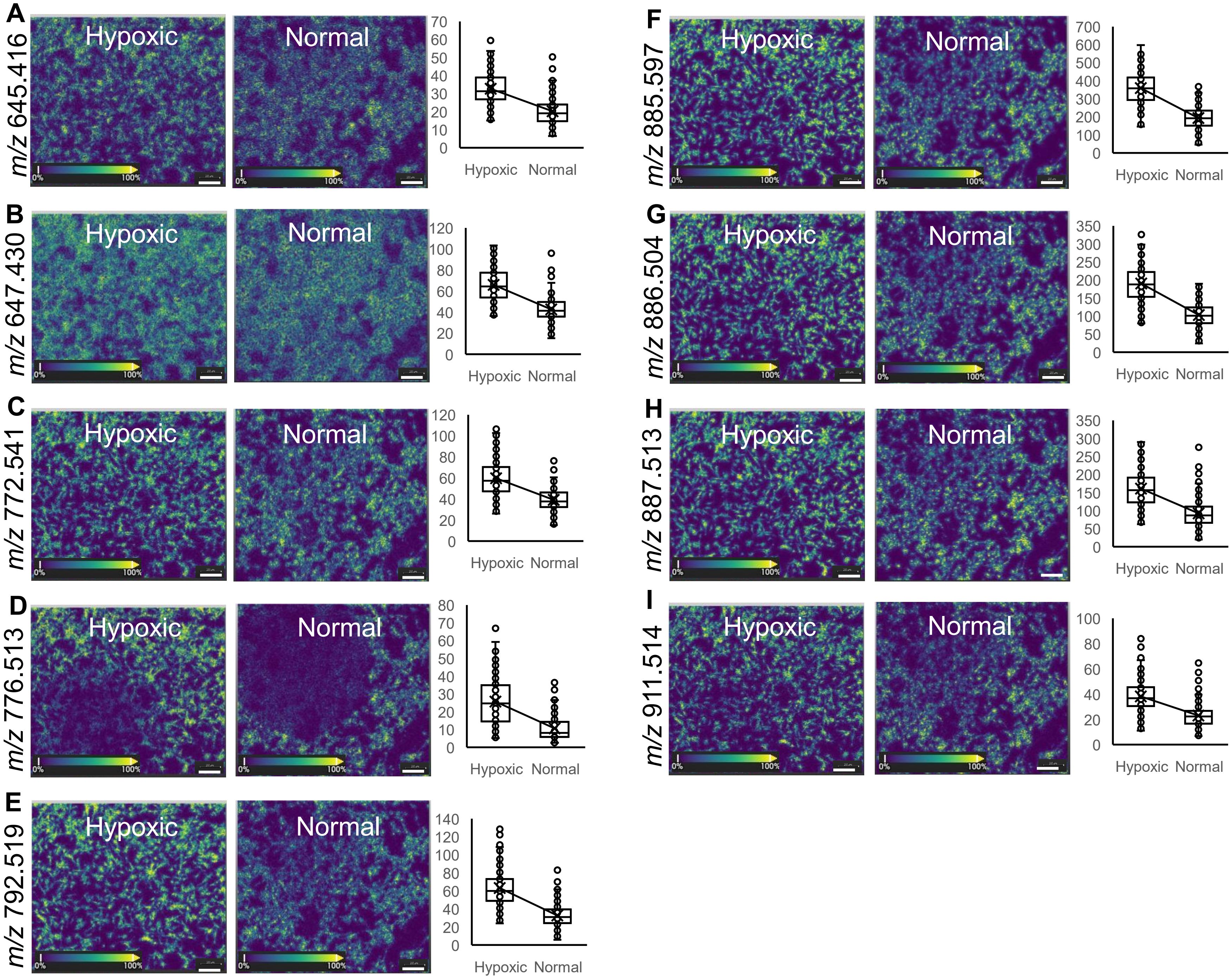
