## Supplementary Figure 2 for "Hypoxia induced lipid alterations in iPSC-derived human glial progenitor cells revealed by Matrix assisted laser desorption ionization mass spectrometry based cellular fingerprinting"

### Supplementary Fig. 2

A

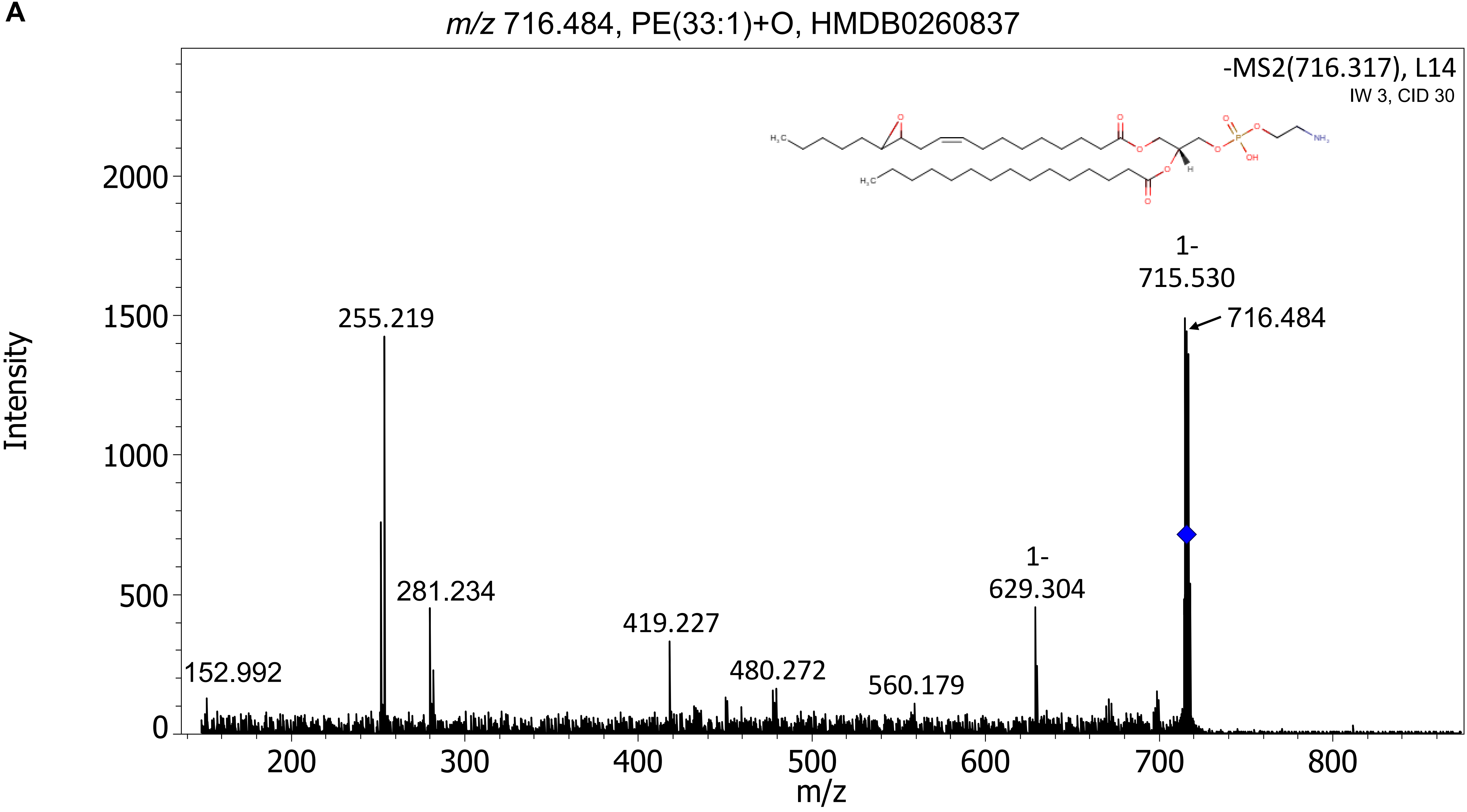

m/z 716.484, PE(33:1)+O, HMDB0260837
-MS2(716.317), L14
IW 3, CID 30
Intensity
m/z
255.219
281.234
419.227
480.272
560.179
152.992
1-
629.304
1-
715.530
716.484

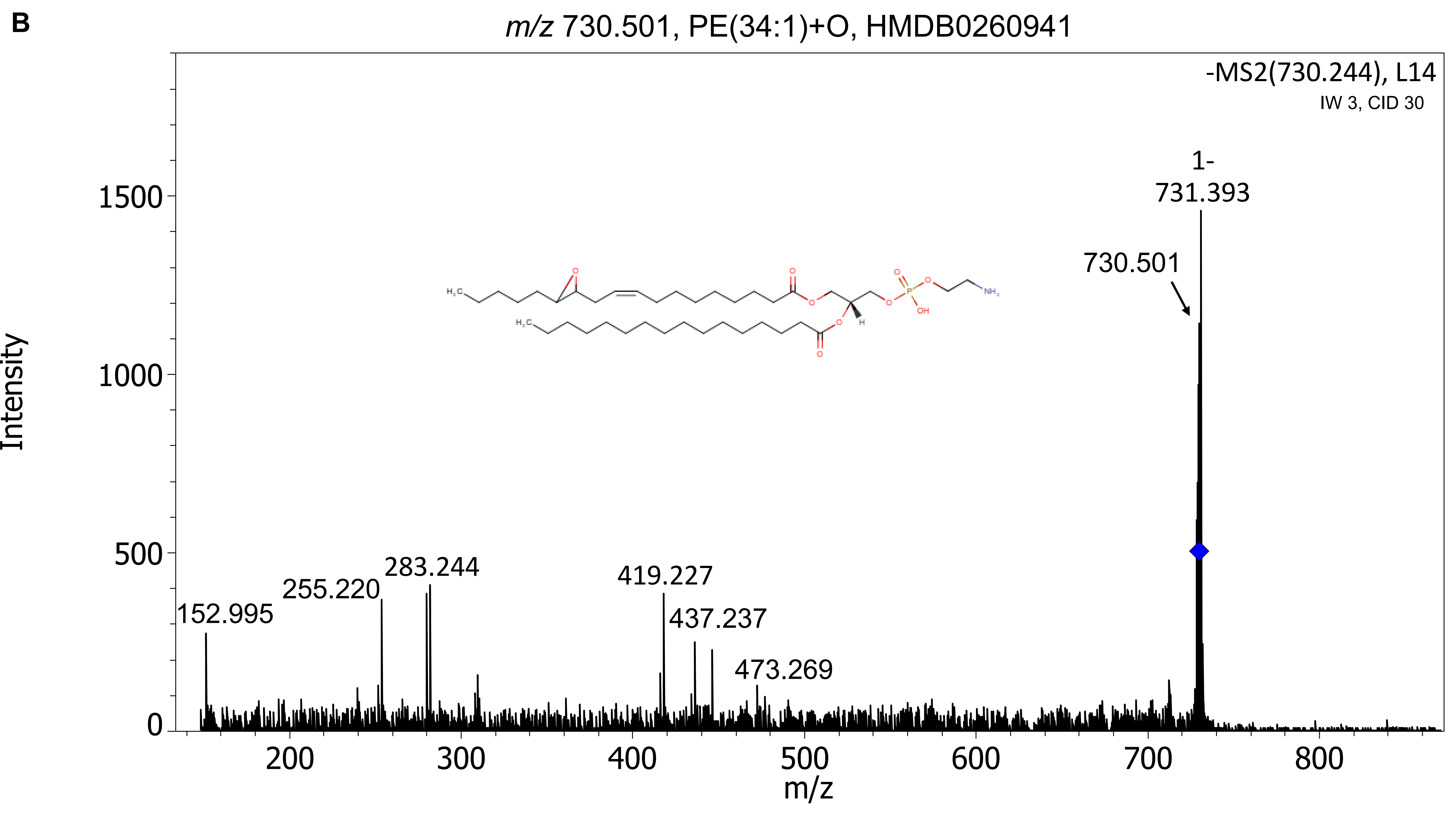

B
m/z 730.501, PE(34:1)+O, HMDB0260941
-MS2(730.244), L14
IW 3, CID 30
1-
731.393
730.501
152.995
255.220
283.244
419.227
437.237
473.269
Intensity
m/z
0
500
1000
1500
200
300
400
500
600
700
800

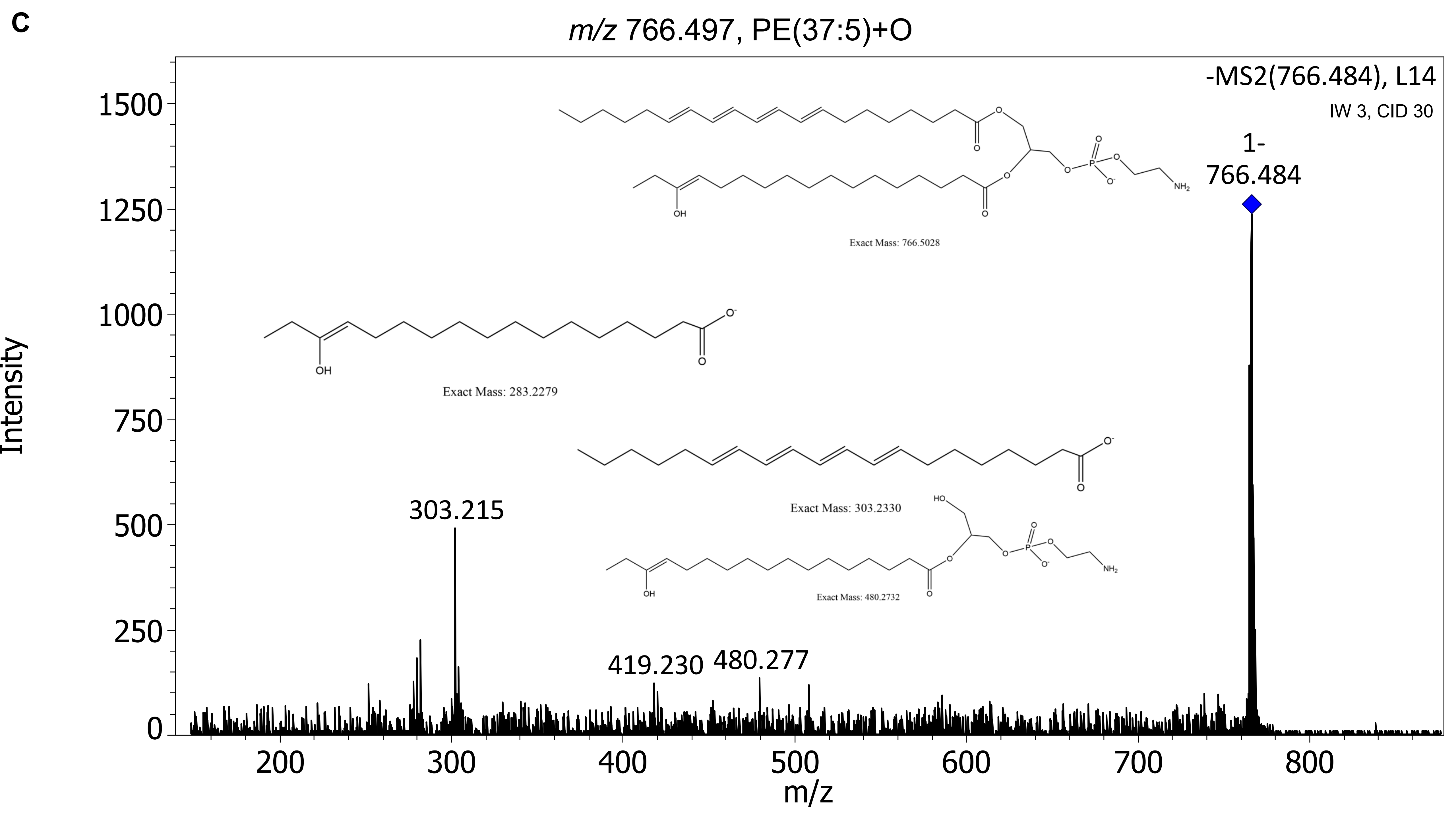
c
m/z 766.497, PE(37:5)+O
-MS2(766.484), L14
IW 3, CID 30
1-
766.484
Exact Mass: 766.5028
Exact Mass: 283.2279
Exact Mass: 303.2330
Exact Mass: 480.2732
303.215
419.230
480.277
Intensity
m/z

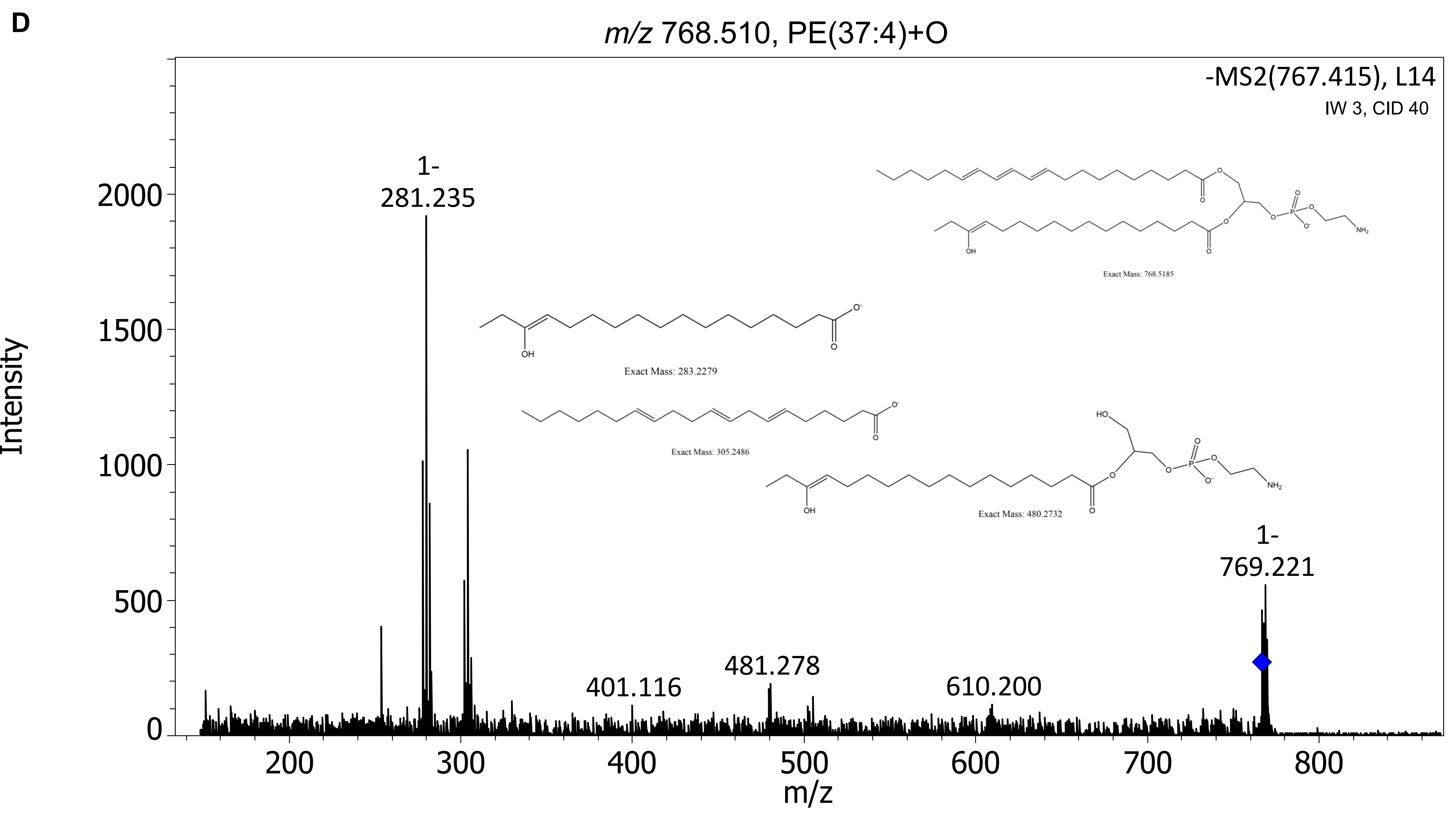
D
m/z 768.510, PE(37:4)+O
-MS2(767.415), L14
IW 3, CID 40
Exact Mass: 768.5185
Exact Mass: 283.2279
Exact Mass: 305.2486
Exact Mass: 480.2732
1- 281.235
1- 769.221
401.116
481.278
610.200
Intensity
m/z

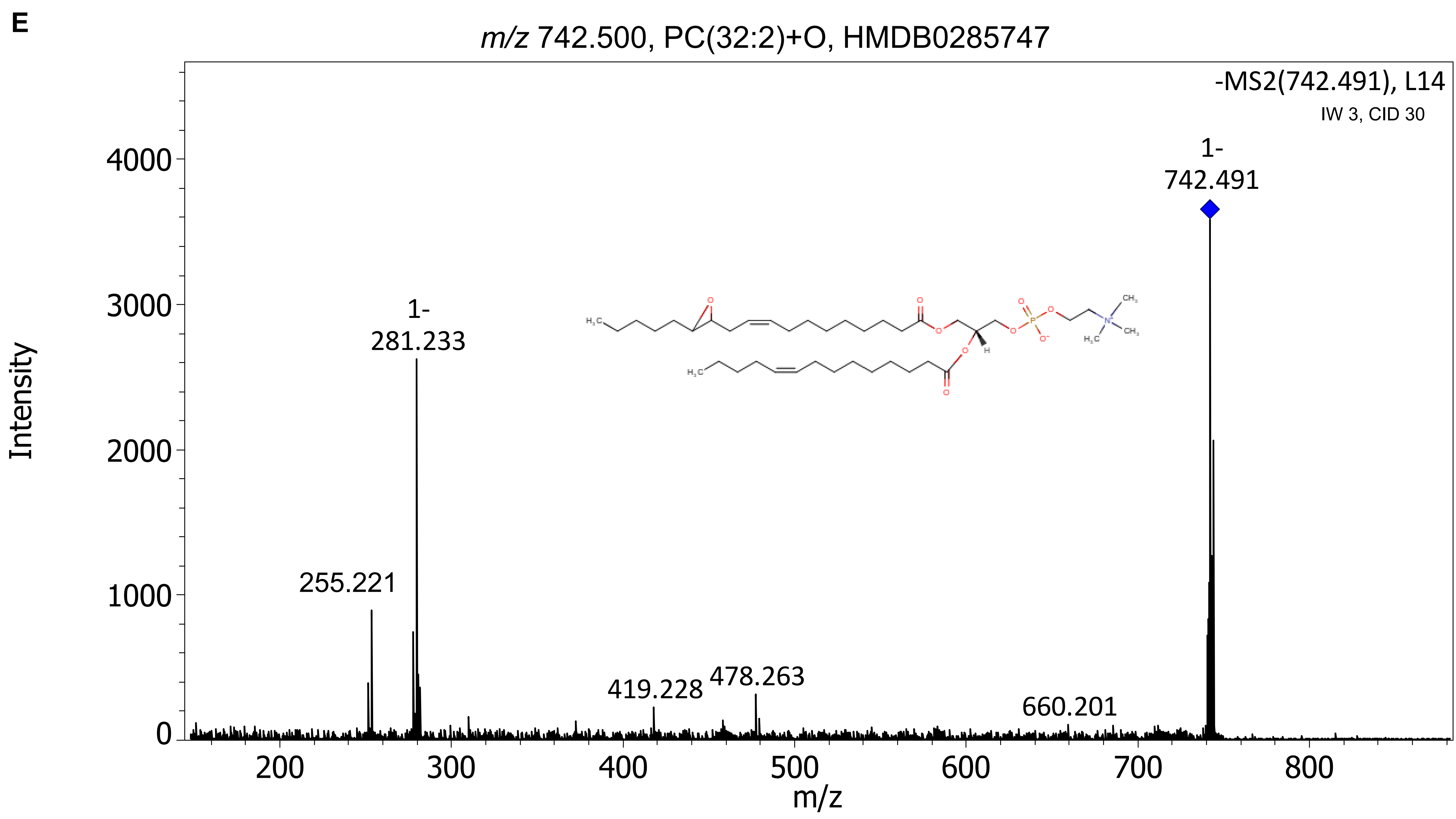
E
m/z 742.500, PC(32:2)+O, HMDB0285747
-MS2(742.491), L14
IW 3, CID 30
1-
742.491
1-
281.233
255.221
419.228
478.263
660.201
Intensity
m/z
0
1000
2000
3000
4000
200
300
400
500
600
700
800

F

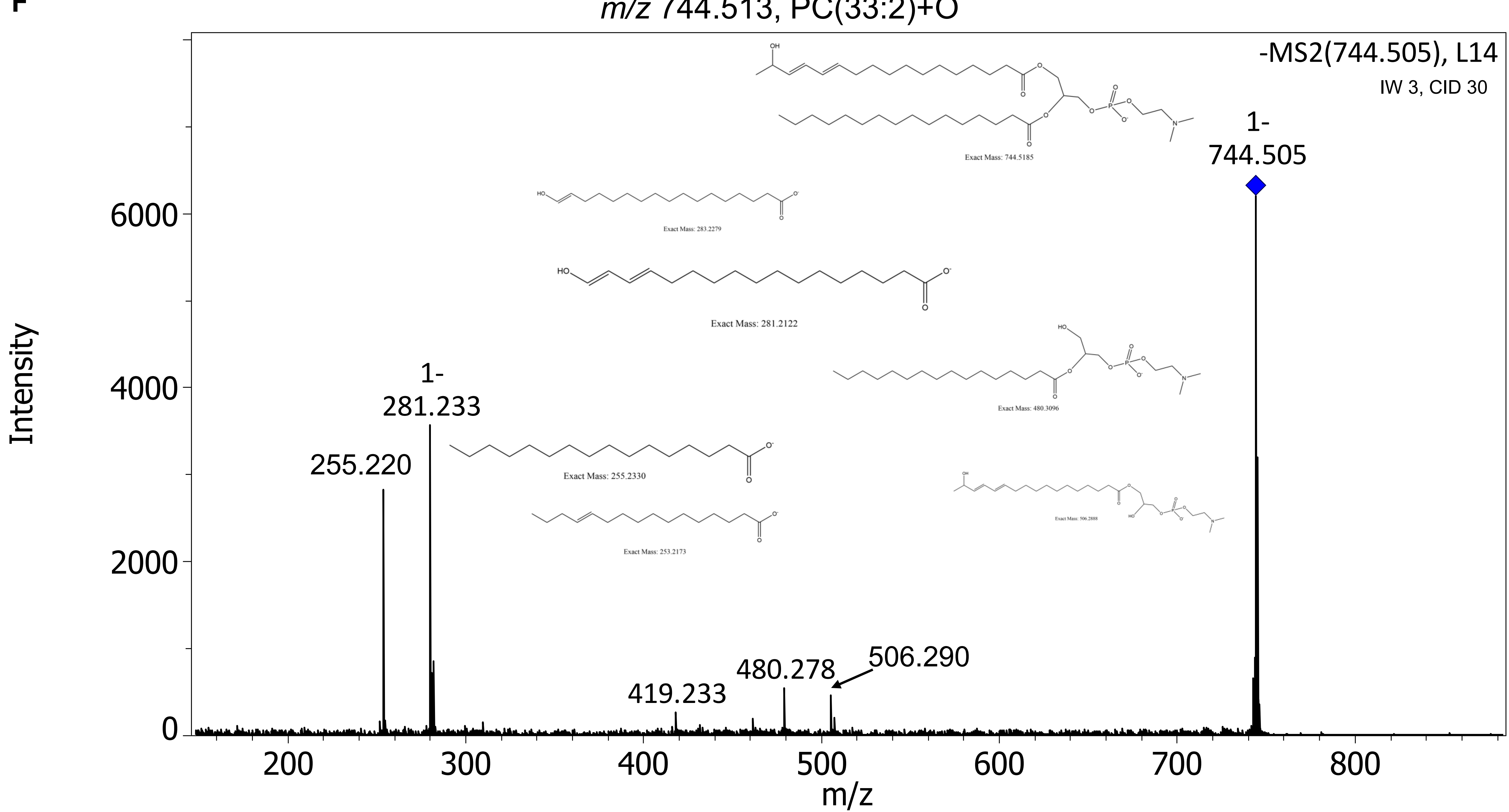

m/z 744.513, PC(33:2)+O
-MS2(744.505), L14
IW 3, CID 30
Exact Mass: 744.5185
1-
744.505
Exact Mass: 283.2279
Exact Mass: 281.2122
Exact Mass: 480.3096
1-
281.233
255.220
Exact Mass: 255.2330
Exact Mass: 506.2888
Exact Mass: 253.2173
419.233
480.278
506.290
Intensity
m/z

G

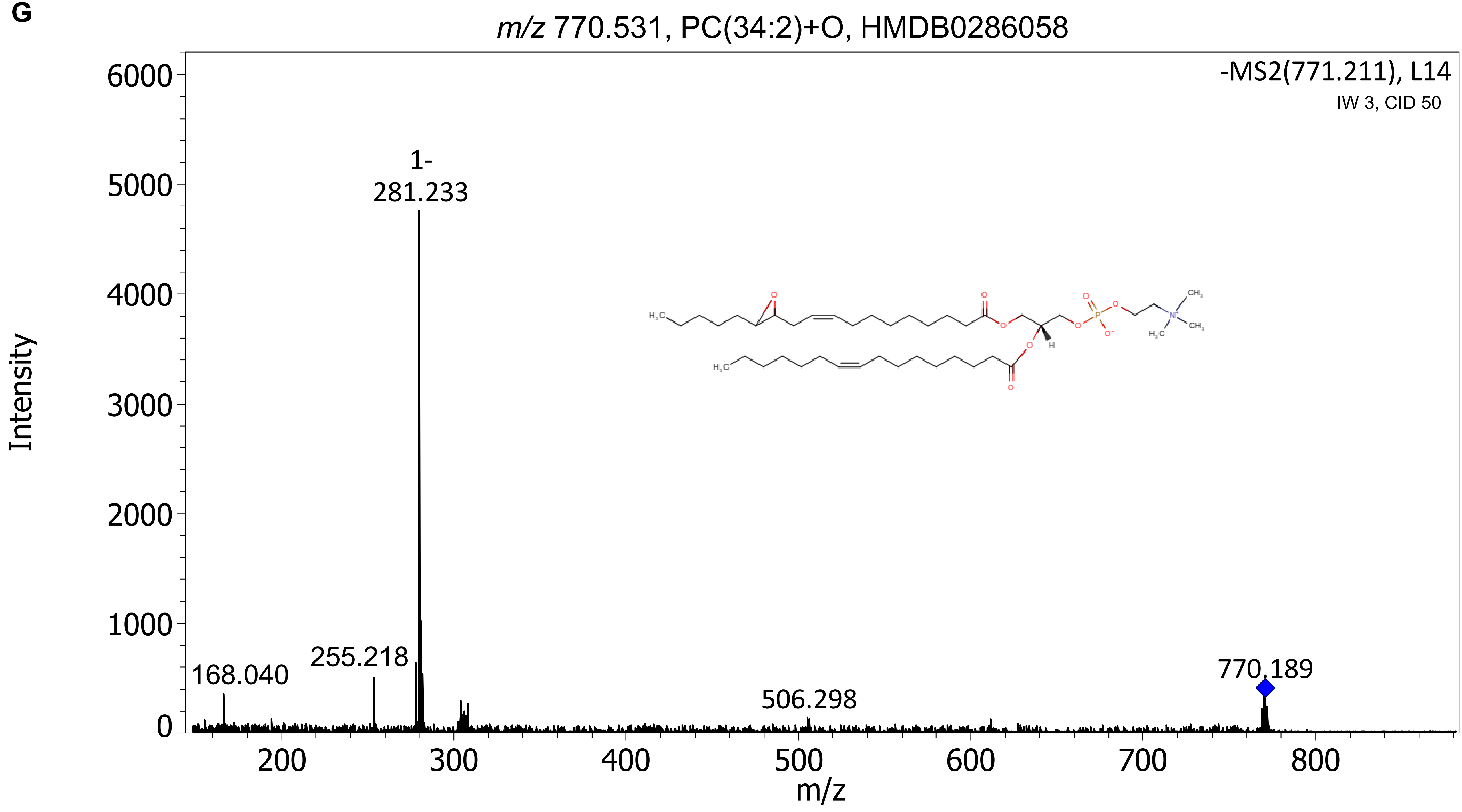
m/z 770.531, PC(34:2)+O, HMDB0286058
-MS2(771.211), L14
IW 3, CID 50
Intensity
m/z
1-
281.233
255.218
168.040
506.298
770.189
6000
5000
4000
3000
2000
1000
0
200
300
400
500
600
700
800

H

*m/z* 794.534, PC(34:1)+O, HMDB0285954

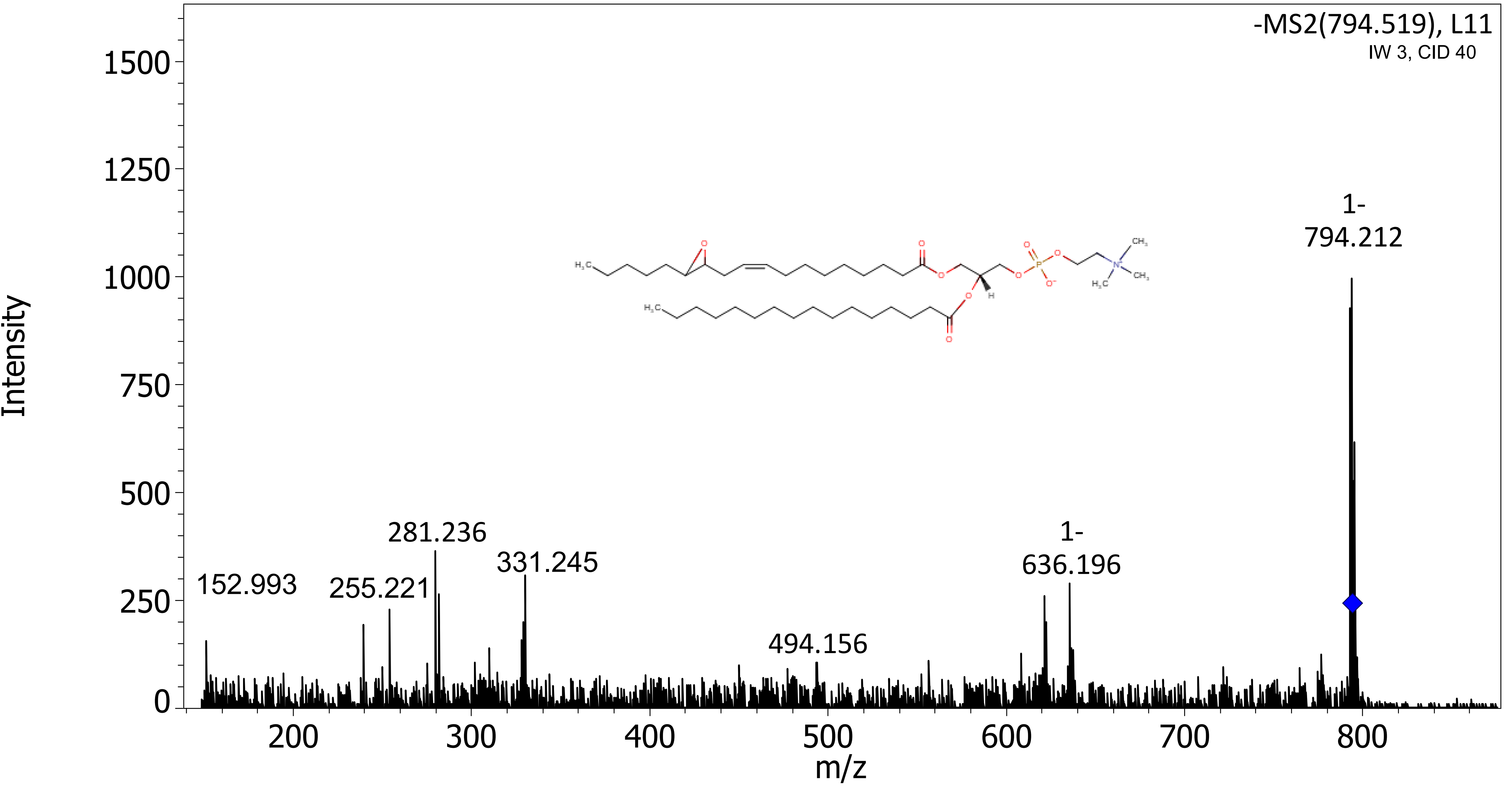

I

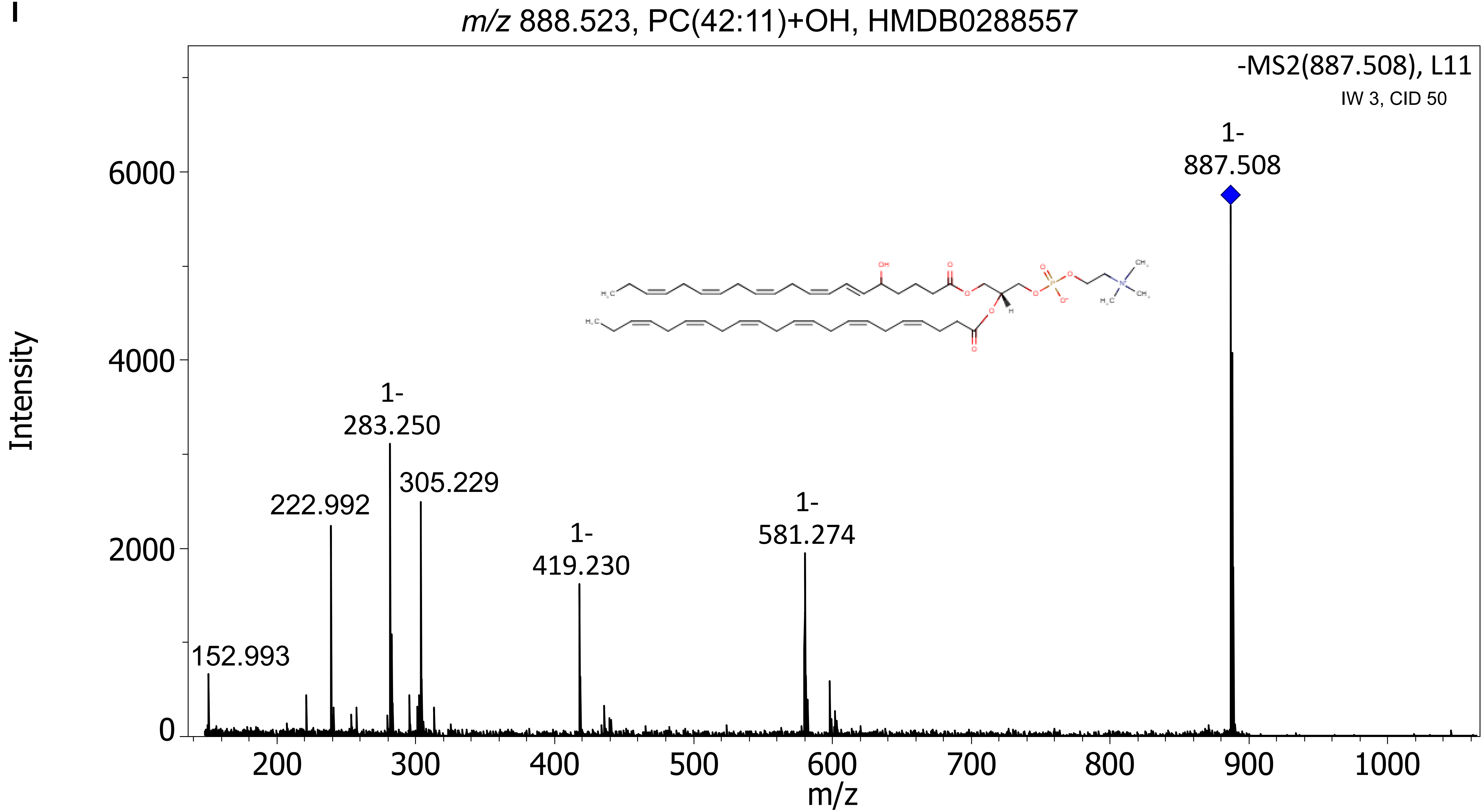

m/z 888.523, PC(42:11)+OH, HMDB0288557
-MS2(887.508), L11
IW 3, CID 50
1-
887.508
1-
283.250
305.229
222.992
1-
419.230
1-
581.274
152.993
Intensity
0
2000
4000
6000
200
300
400
500
600
700
800
900
1000
m/z

J

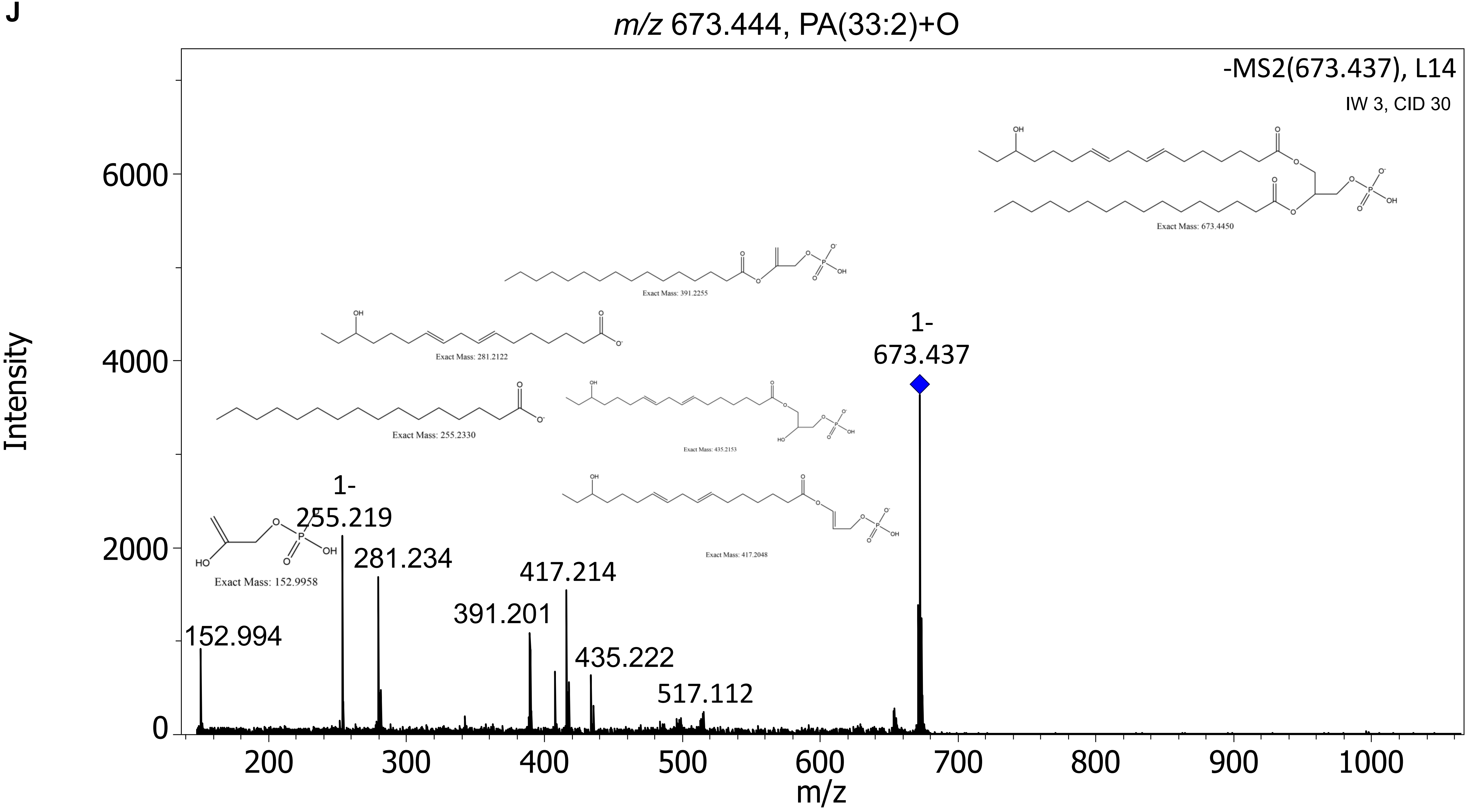
m/z 673.444, PA(33:2)+O
-MS2(673.437), L14
IW 3, CID 30
Exact Mass: 673.4450
Exact Mass: 391.2255
Exact Mass: 281.2122
Exact Mass: 255.2330
Exact Mass: 435.2153
Exact Mass: 417.2048
Exact Mass: 152.9958
1-
673.437
1-
255.219
281.234
417.214
391.201
152.994
435.222
517.112
Intensity
m/z
0
2000
4000
6000
200
300
400
500
600
700
800
900
1000

K

*m/z* 699.466, PA(35:3)+OH, HMDB0267921

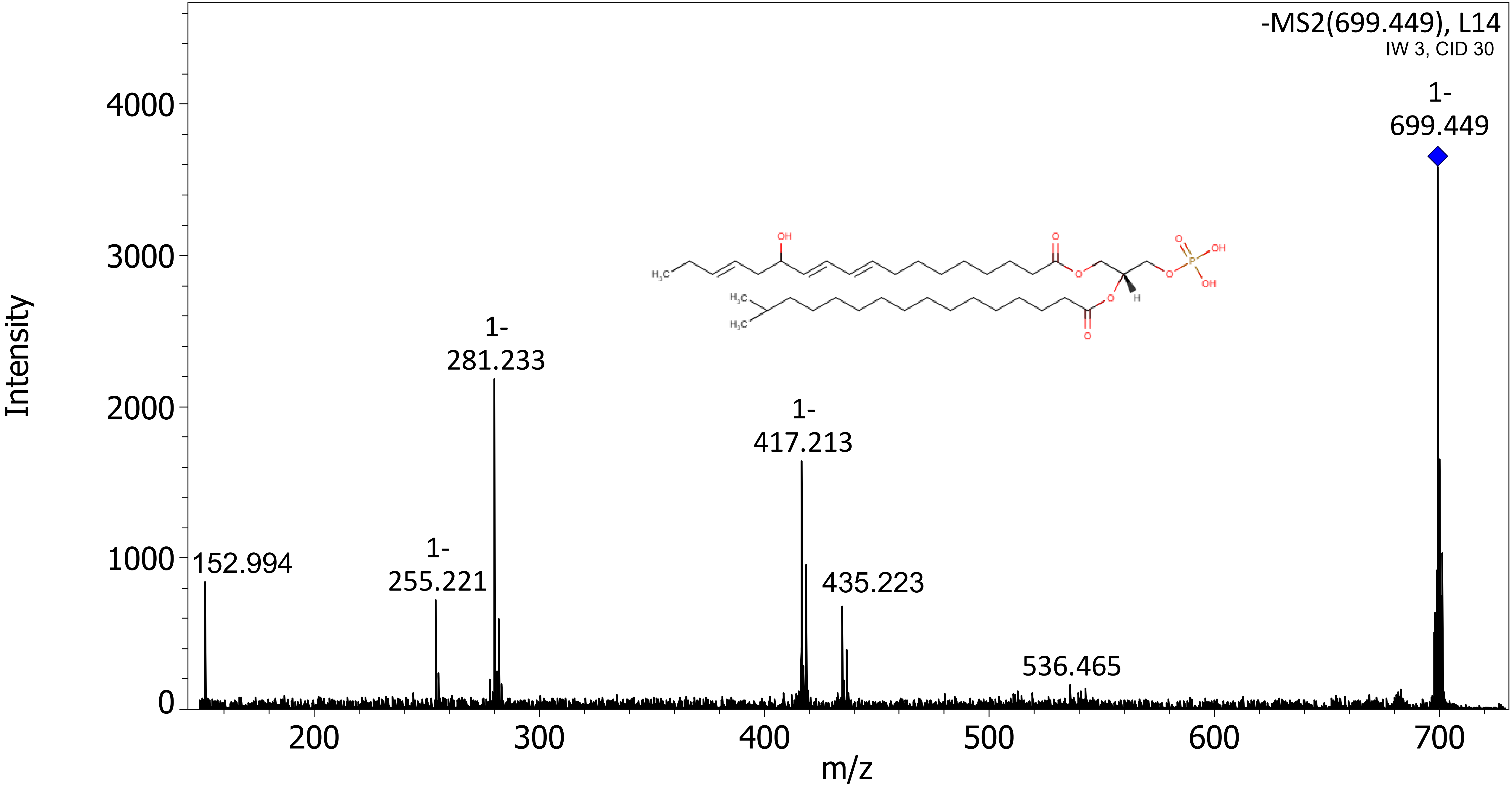

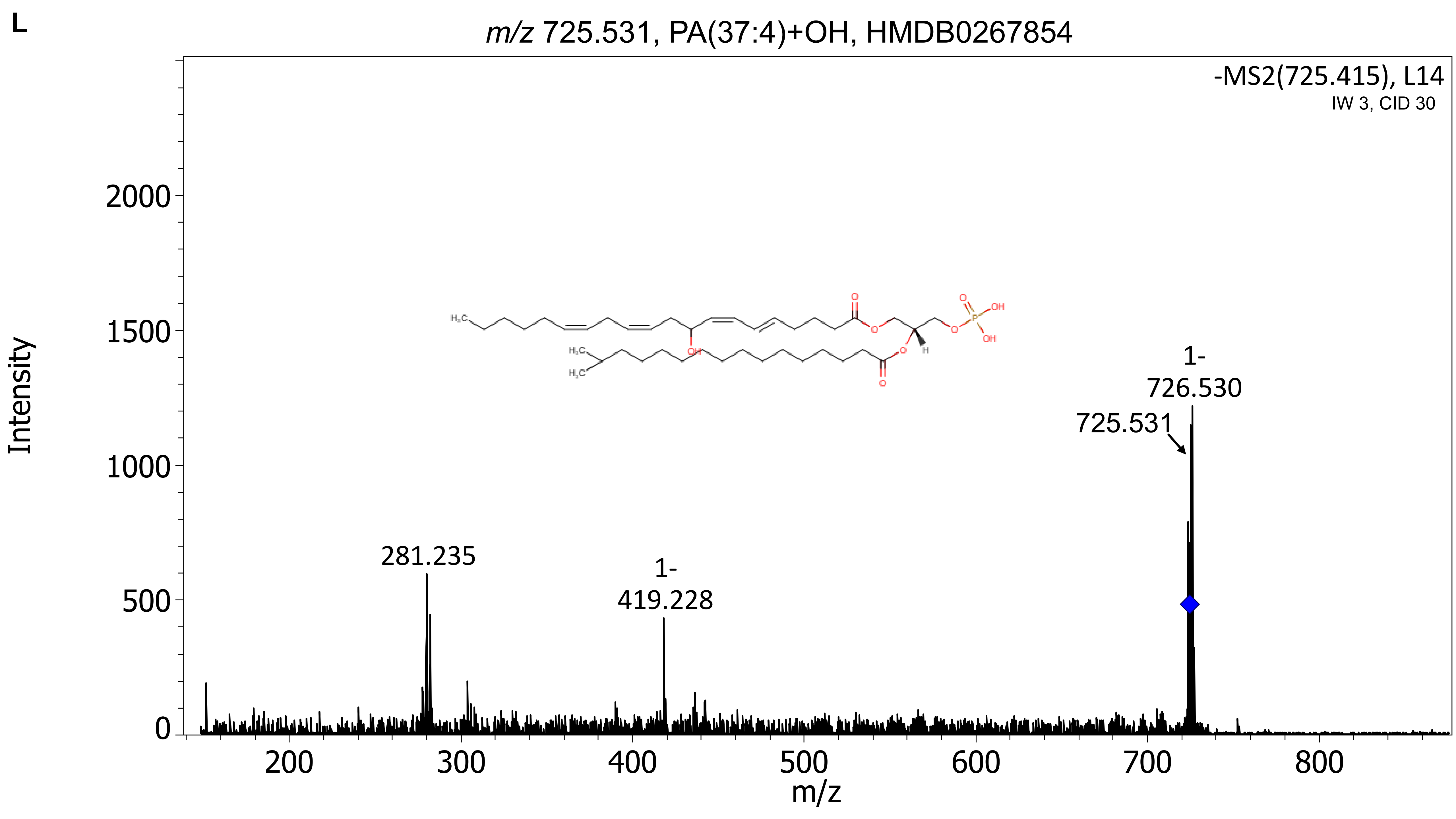
L
m/z 725.531, PA(37:4)+OH, HMDB0267854
-MS2(725.415), L14
IW 3, CID 30
Intensity
m/z
281.235
1-
419.228
725.531
1-
726.530
0
500
1000
1500
2000
200
300
400
500
600
700
800

### MS-MS of Unidentified molecules (not found in database)

**M**

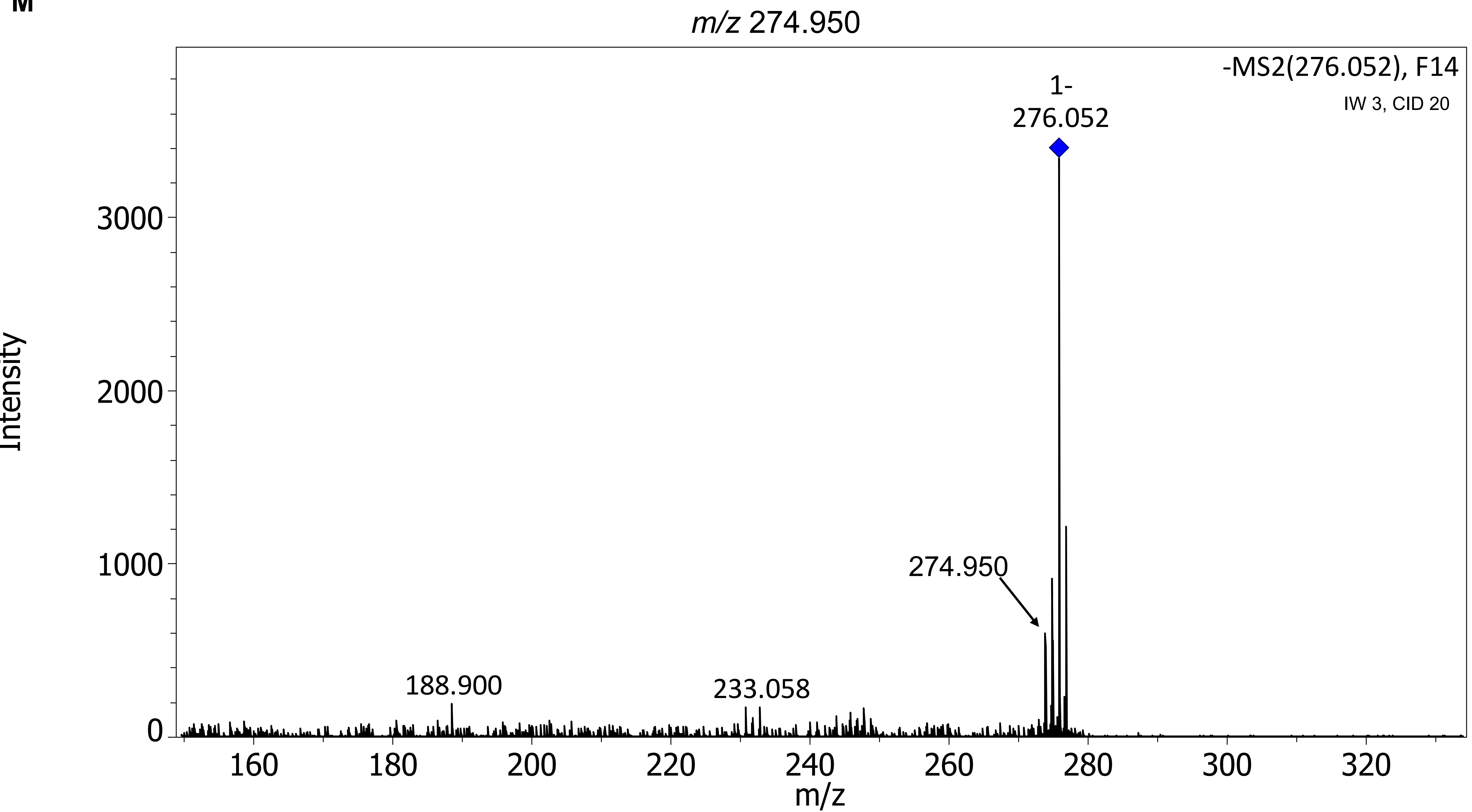

m/z 274.950
-MS2(276.052), F14
IW 3, CID 20
1-
276.052
274.950
188.900
233.058
Intensity
3000
2000
1000
0
160
180
200
220
240
260
280
300
320
m/z

N

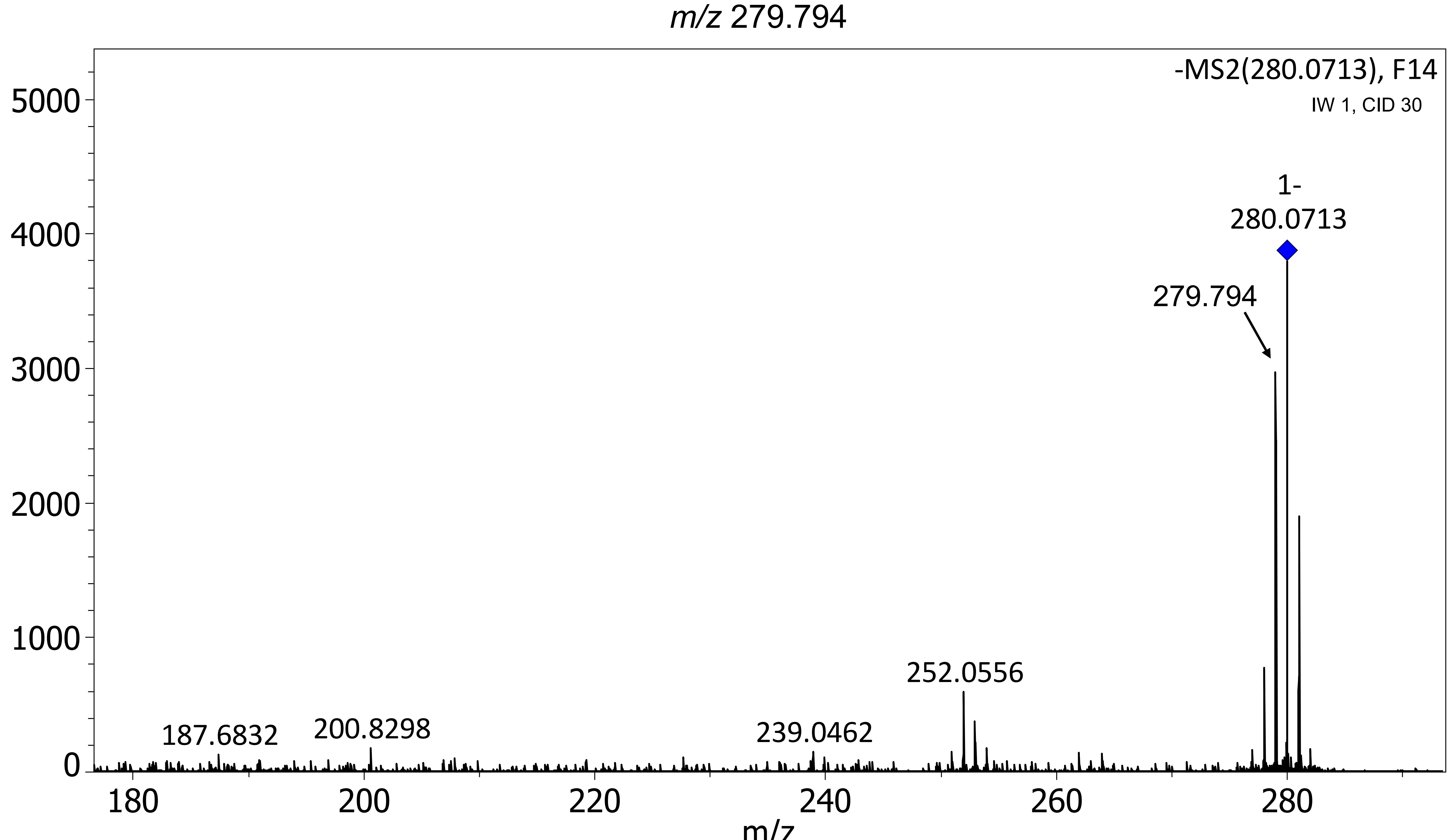
m/z 279.794
-MS2(280.0713), F14
IW 1, CID 30
5000
4000
3000
2000
1000
0
1-
280.0713
279.794
252.0556
239.0462
200.8298
187.6832
180
200
220
240
260
280
m/z

O

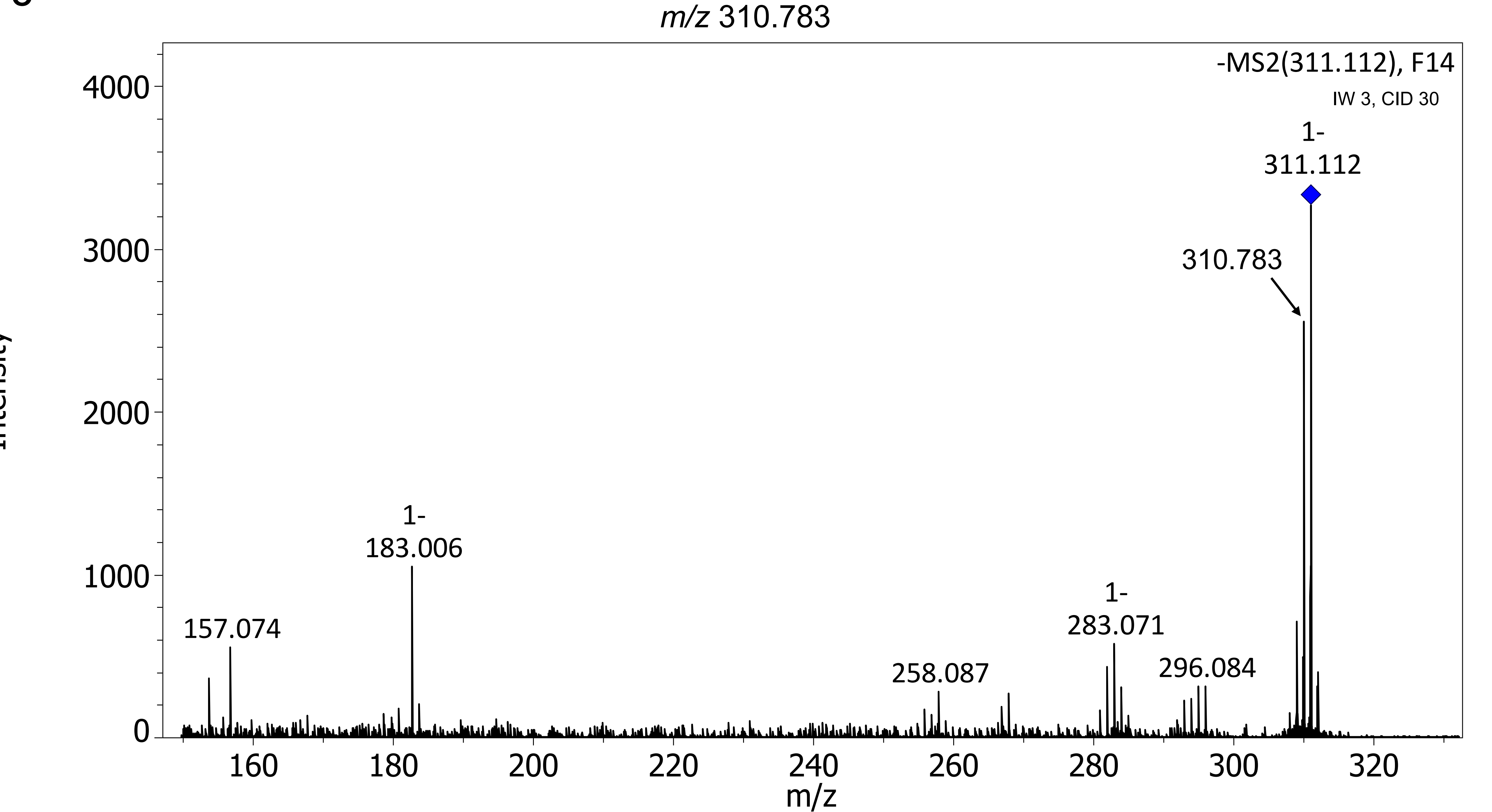

m/z 310.783
-MS2(311.112), F14
IW 3, CID 30
1-
311.112
310.783
1-
183.006
157.074
258.087
1-
283.071
296.084
4000
3000
2000
1000
0
160
180
200
220
240
260
280
300
320
m/z
Intensity

P

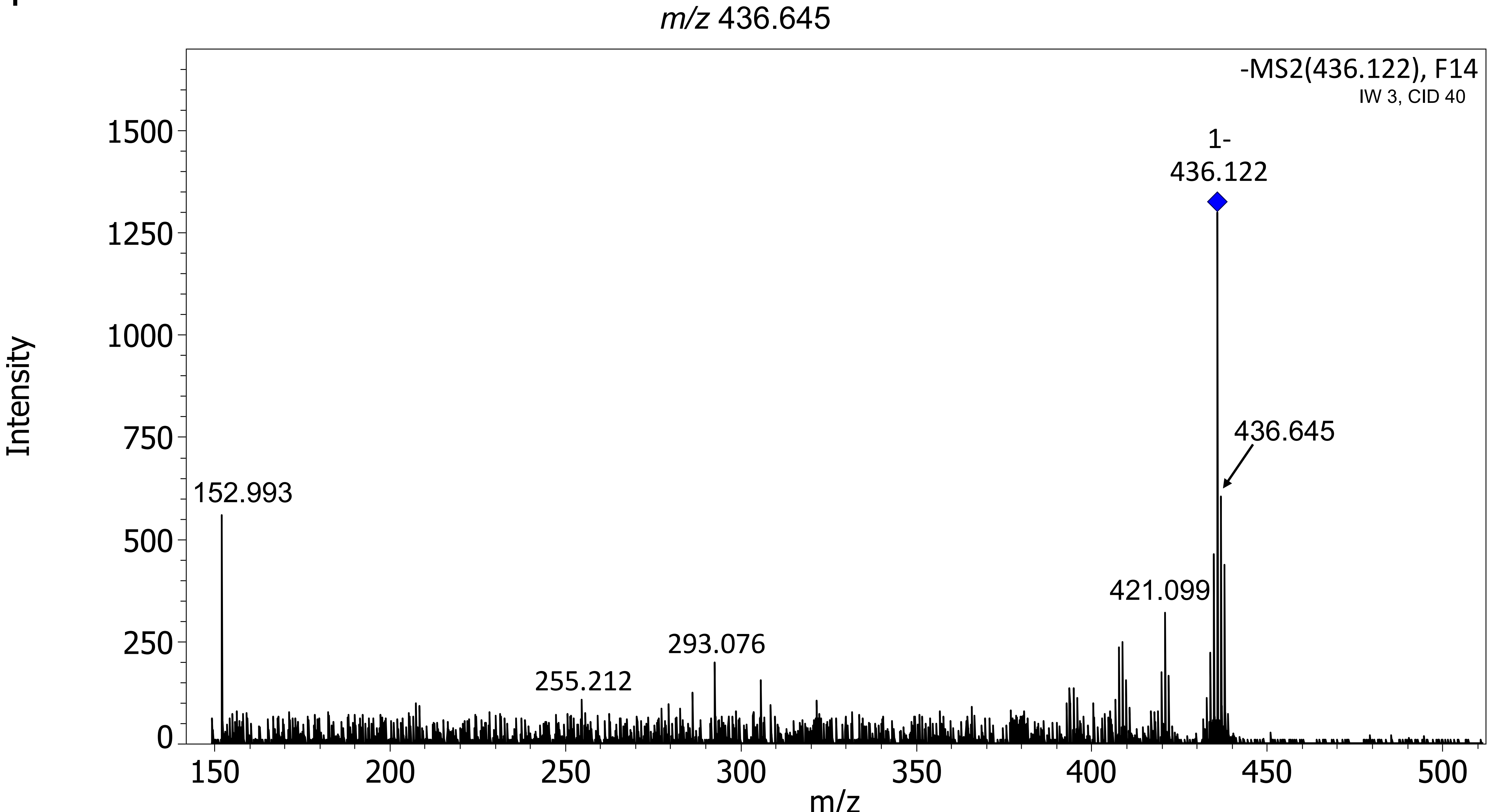
m/z 436.645
-MS2(436.122), F14
IW 3, CID 40
1-
436.122
436.645
152.993
421.099
293.076
255.212
Intensity
m/z
0
250
500
750
1000
1250
1500
150
200
250
300
350
400
450
500

Q

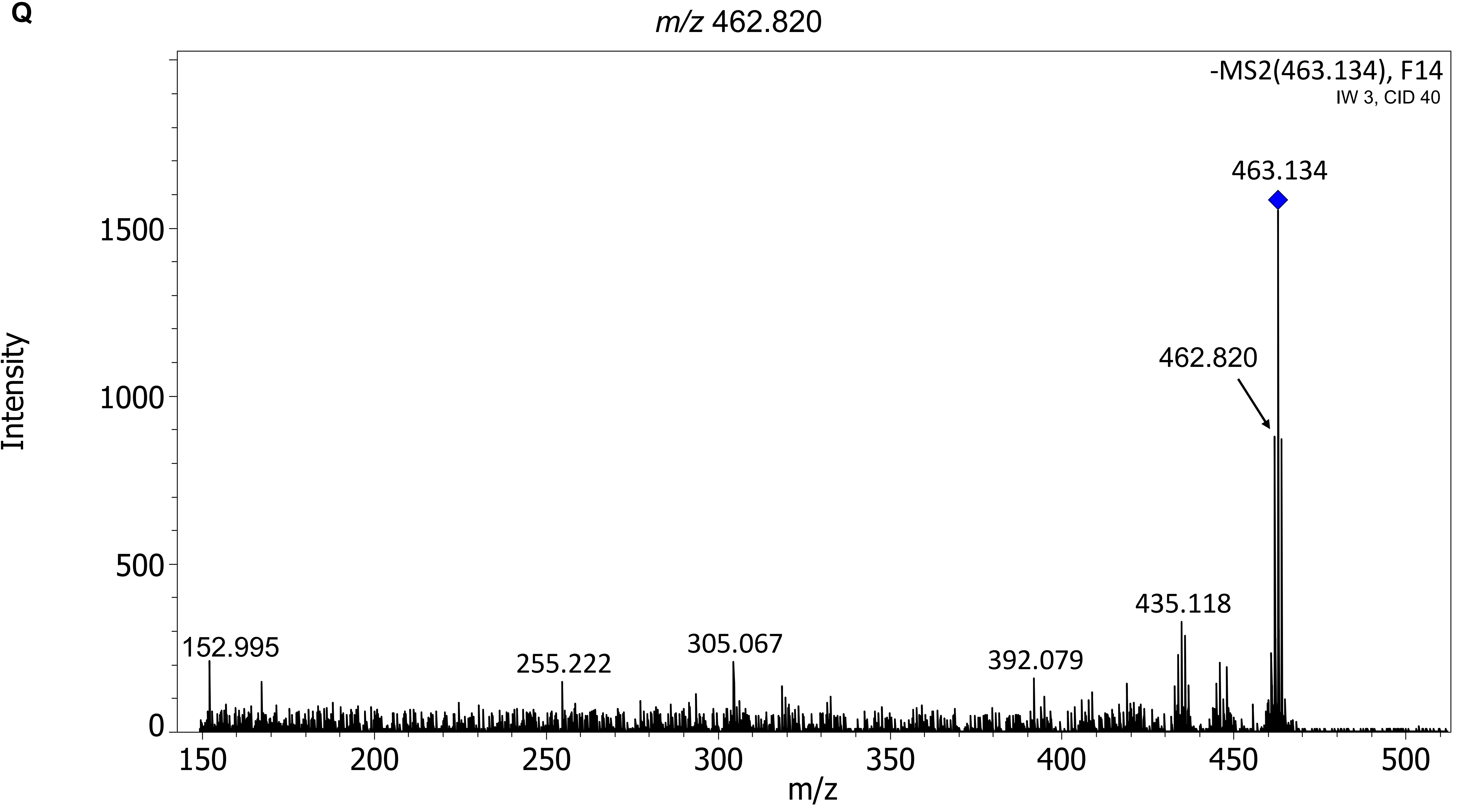

m/z 462.820
-MS2(463.134), F14
IW 3, CID 40
463.134
462.820
435.118
392.079
305.067
255.222
152.995
Intensity
0
500
1000
1500
150
200
250
300
350
400
450
500
m/z

**R**

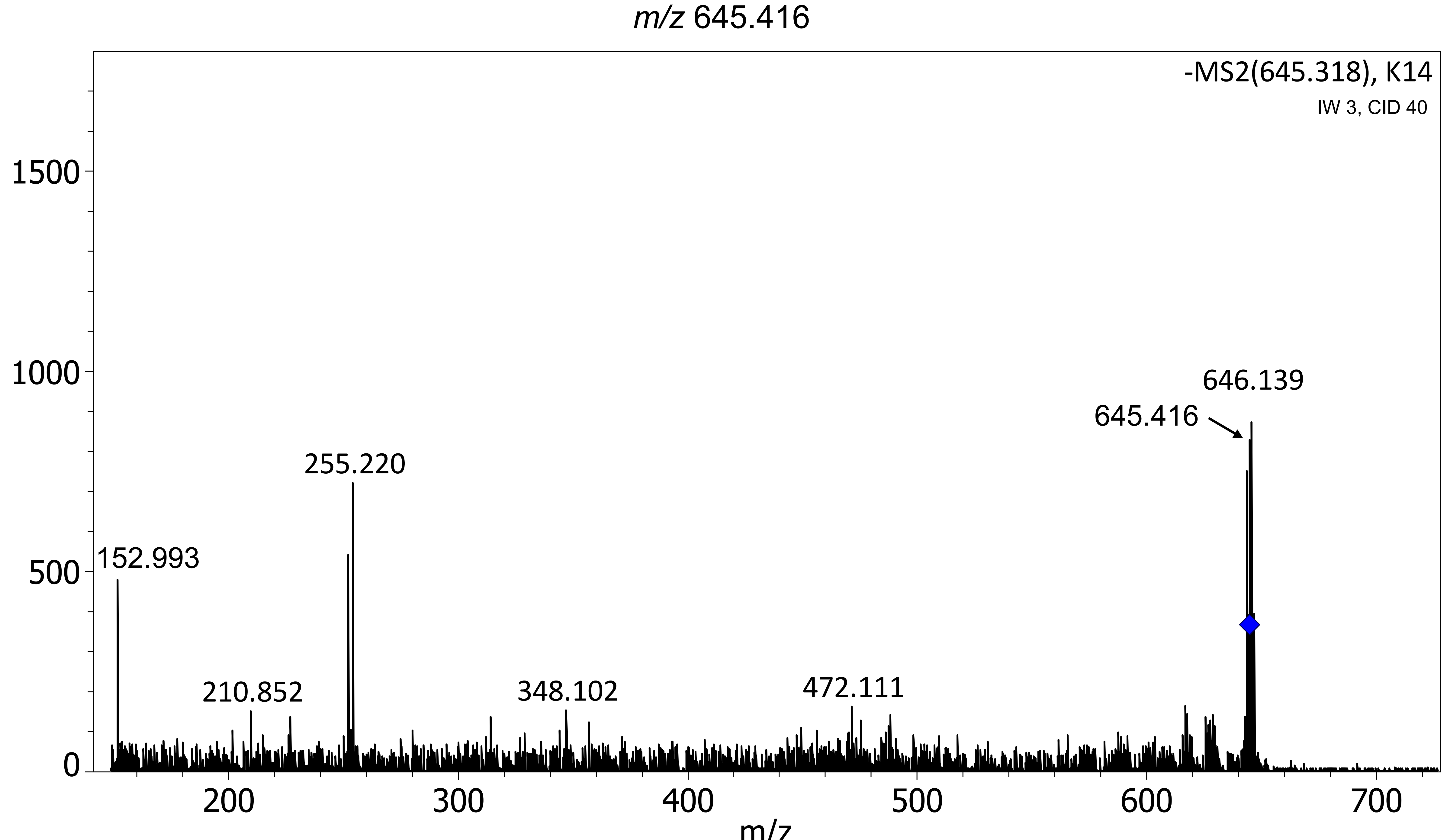

m/z 645.416
-MS2(645.318), K14
IW 3, CID 40
1500
1000
500
0
646.139
645.416
255.220
152.993
210.852
348.102
472.111
200
300
400
500
600
700
m/z

**S**

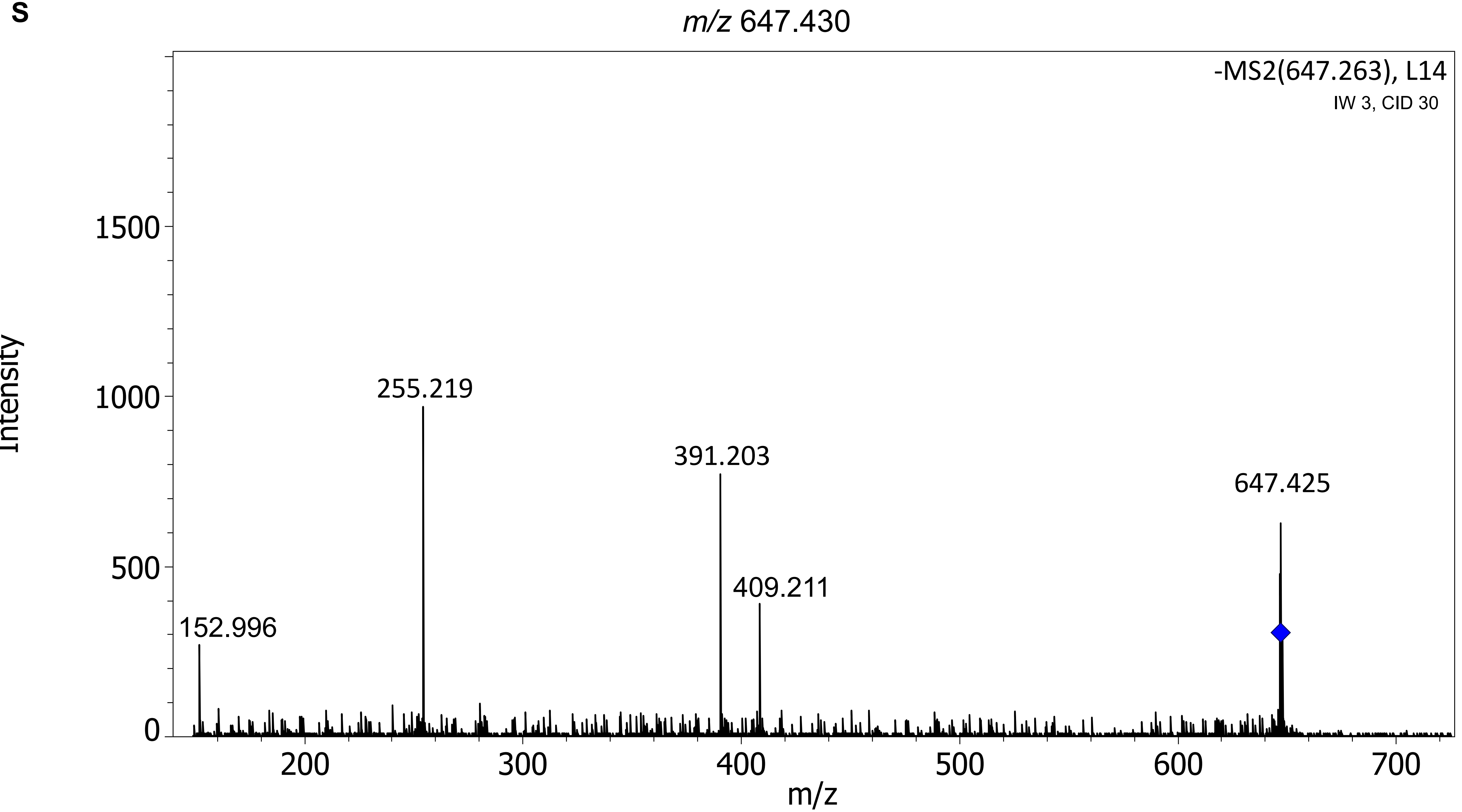

m/z 647.430
-MS2(647.263), L14
IW 3, CID 30
Intensity
m/z
1500
1000
500
0
200
300
400
500
600
700
255.219
391.203
409.211
647.425
152.996

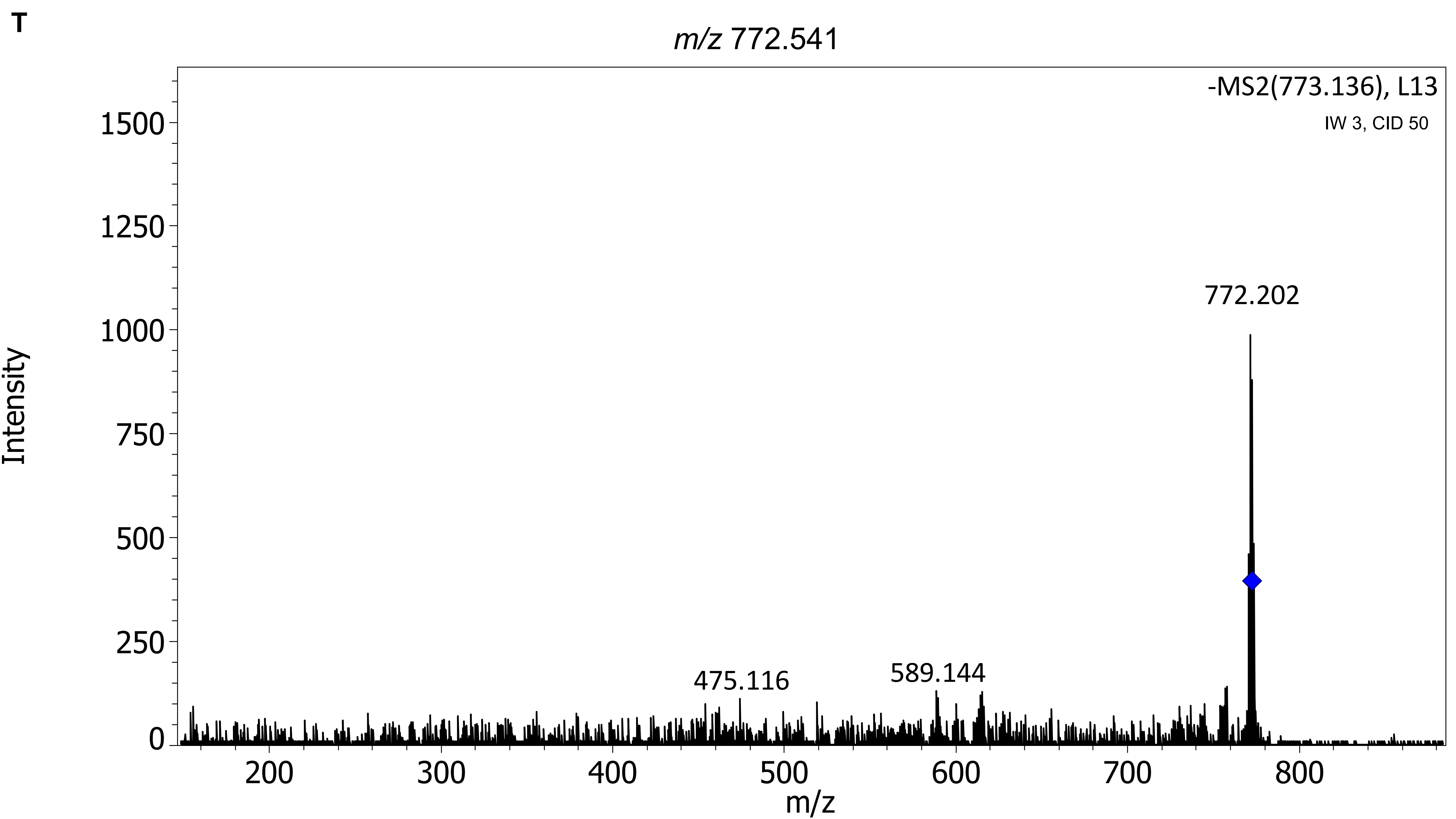
T
m/z 772.541
-MS2(773.136), L13
IW 3, CID 50
772.202
475.116
589.144
Intensity
m/z
0
250
500
750
1000
1250
1500
200
300
400
500
600
700
800

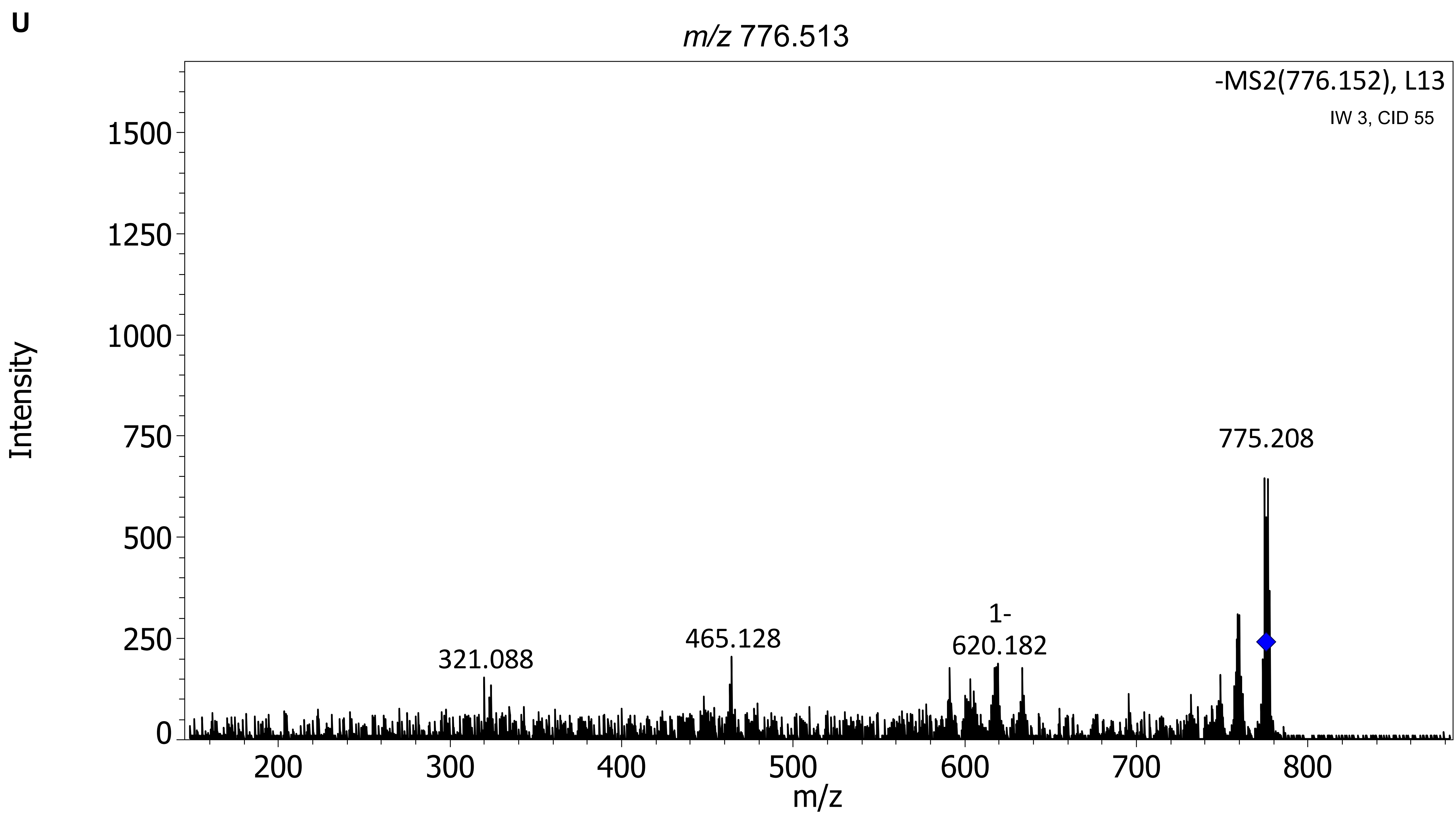
U
m/z 776.513
-MS2(776.152), L13
IW 3, CID 55
Intensity
m/z
321.088
465.128
1-
620.182
775.208
0
250
500
750
1000
1250
1500
200
300
400
500
600
700
800

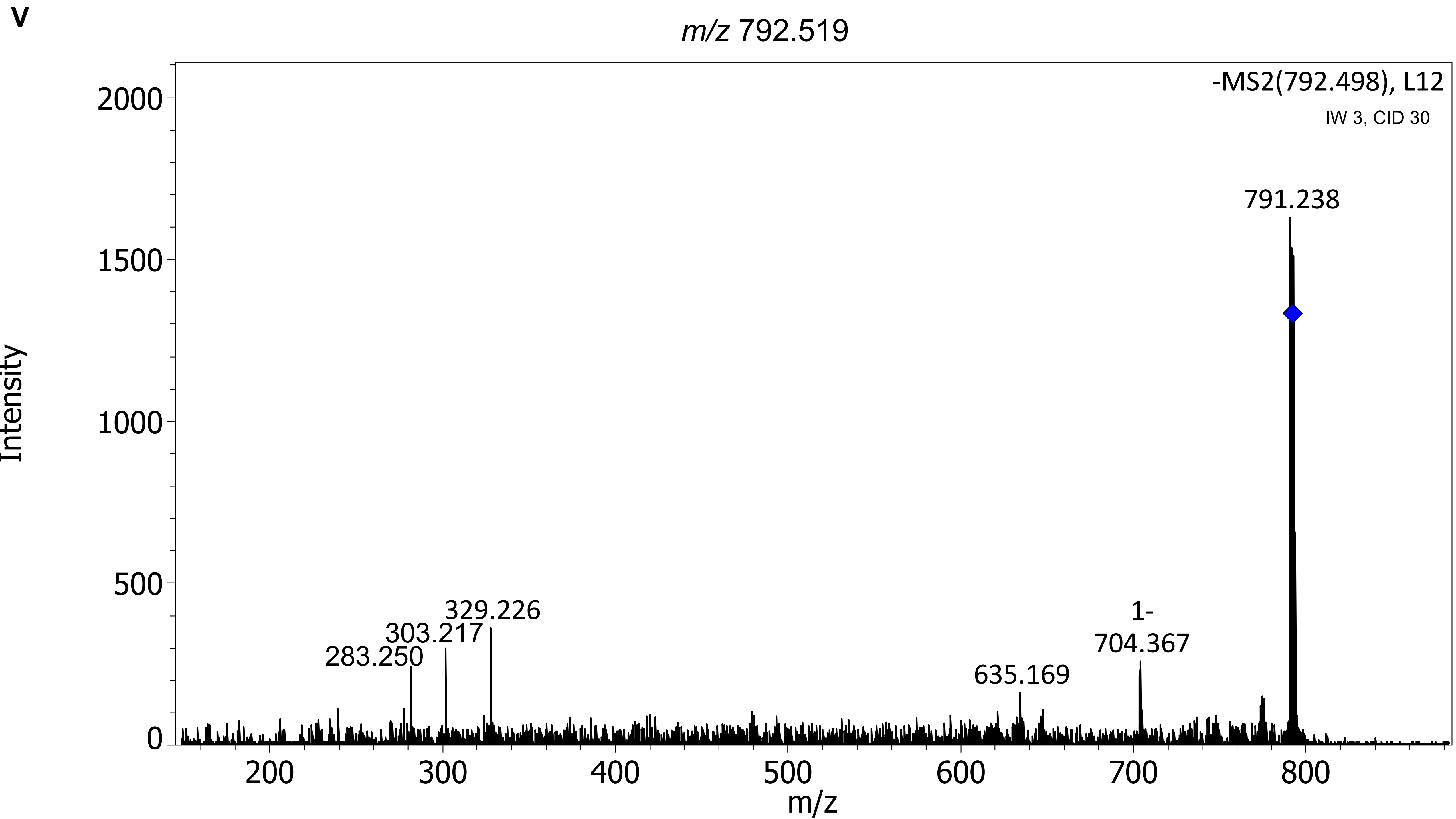
V
m/z 792.519
-MS2(792.498), L12
IW 3, CID 30
791.238
329.226
303.217
283.250
635.169
1-
704.367
Intensity
2000
1500
1000
500
0
200
300
400
500
600
700
800
m/z

**W**

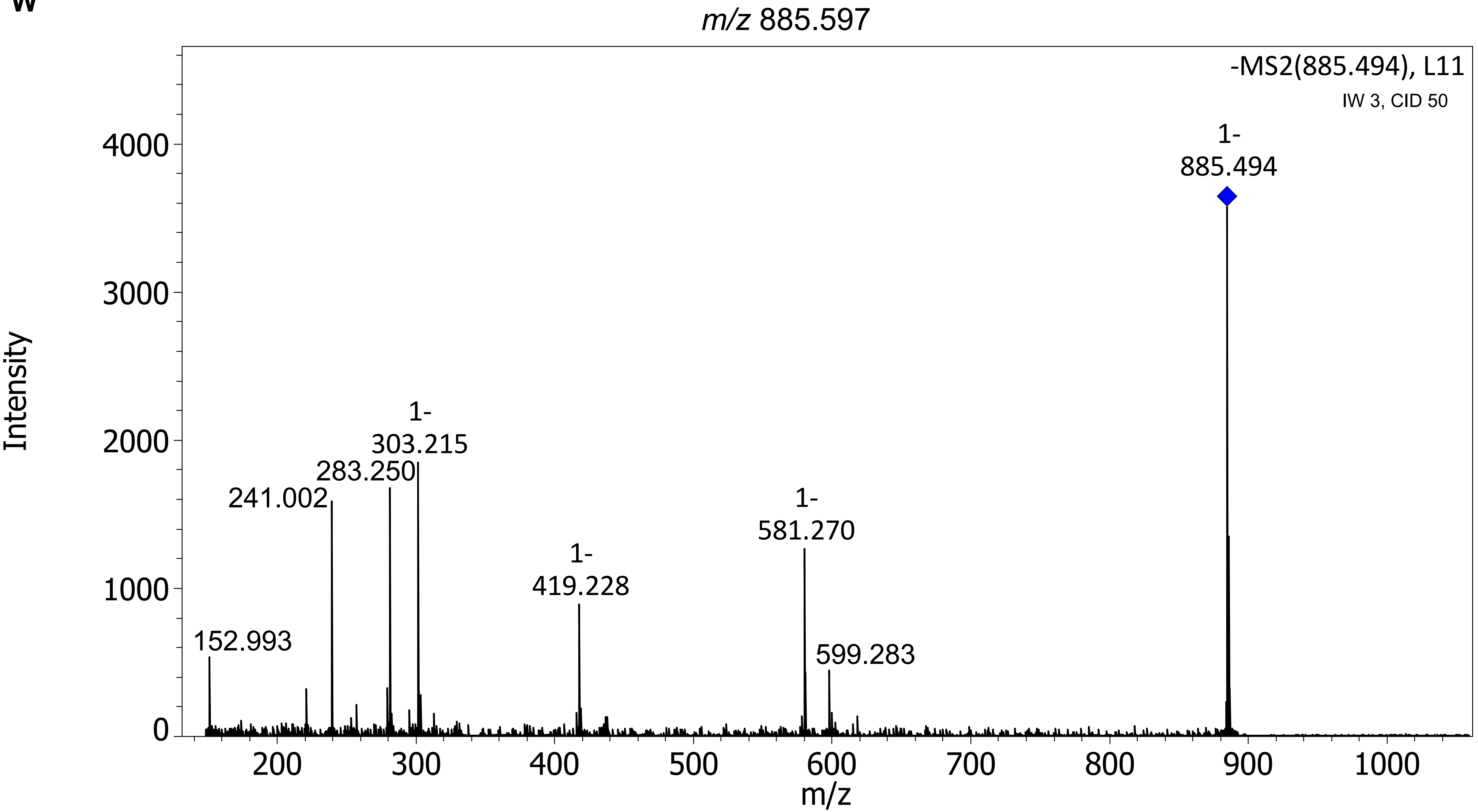

m/z 885.597
-MS2(885.494), L11
IW 3, CID 50
1-
885.494
1-
303.215
283.250
241.002
1-
581.270
1-
419.228
152.993
599.283
Intensity
0
1000
2000
3000
4000
200
300
400
500
600
700
800
900
1000
m/z

**X**

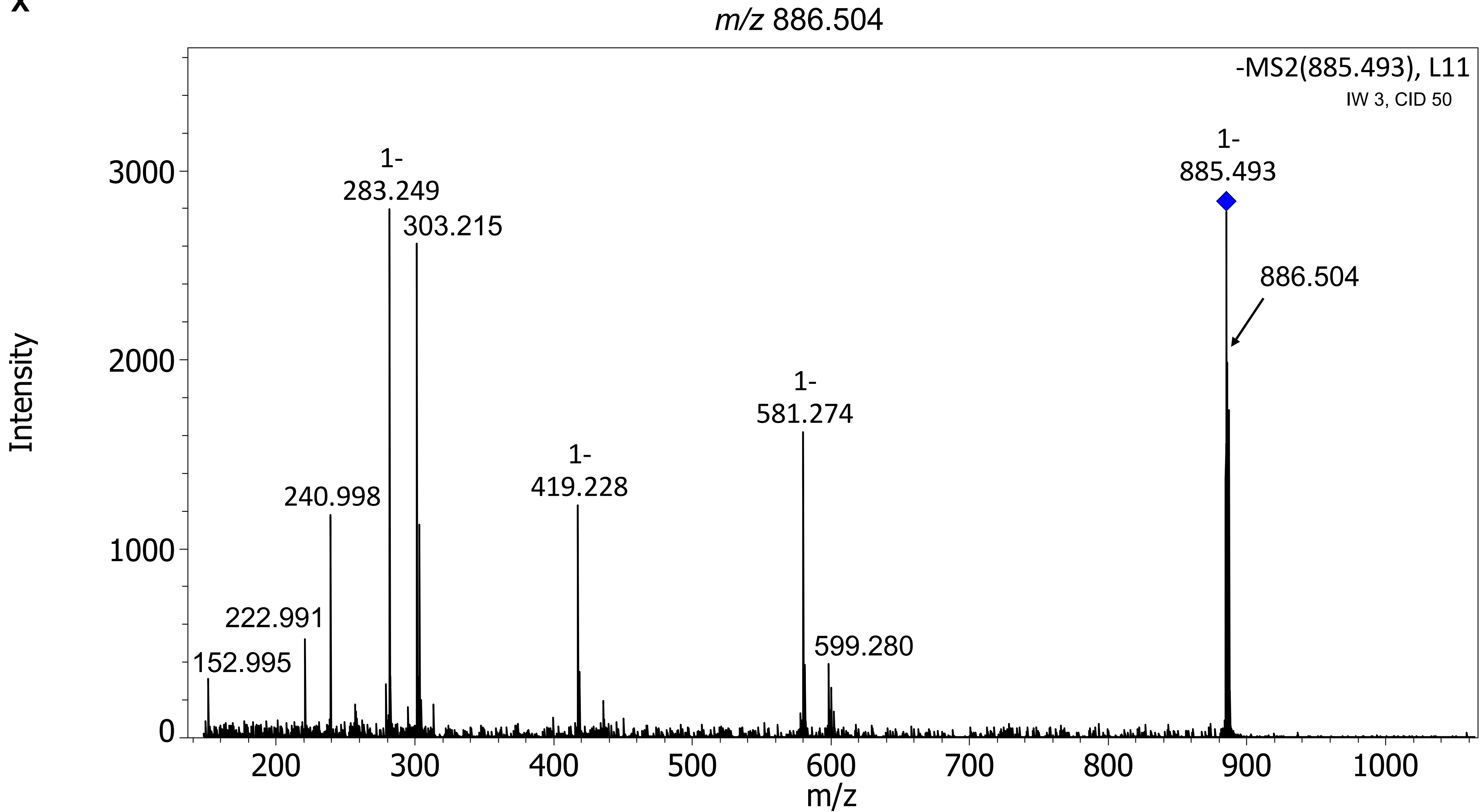
m/z 886.504
-MS2(885.493), L11
IW 3, CID 50
1-
885.493
886.504
1-
283.249
303.215
240.998
222.991
152.995
1-
419.228
1-
581.274
599.280
Intensity
m/z
0
1000
2000
3000
200
300
400
500
600
700
800
900
1000

**Y**

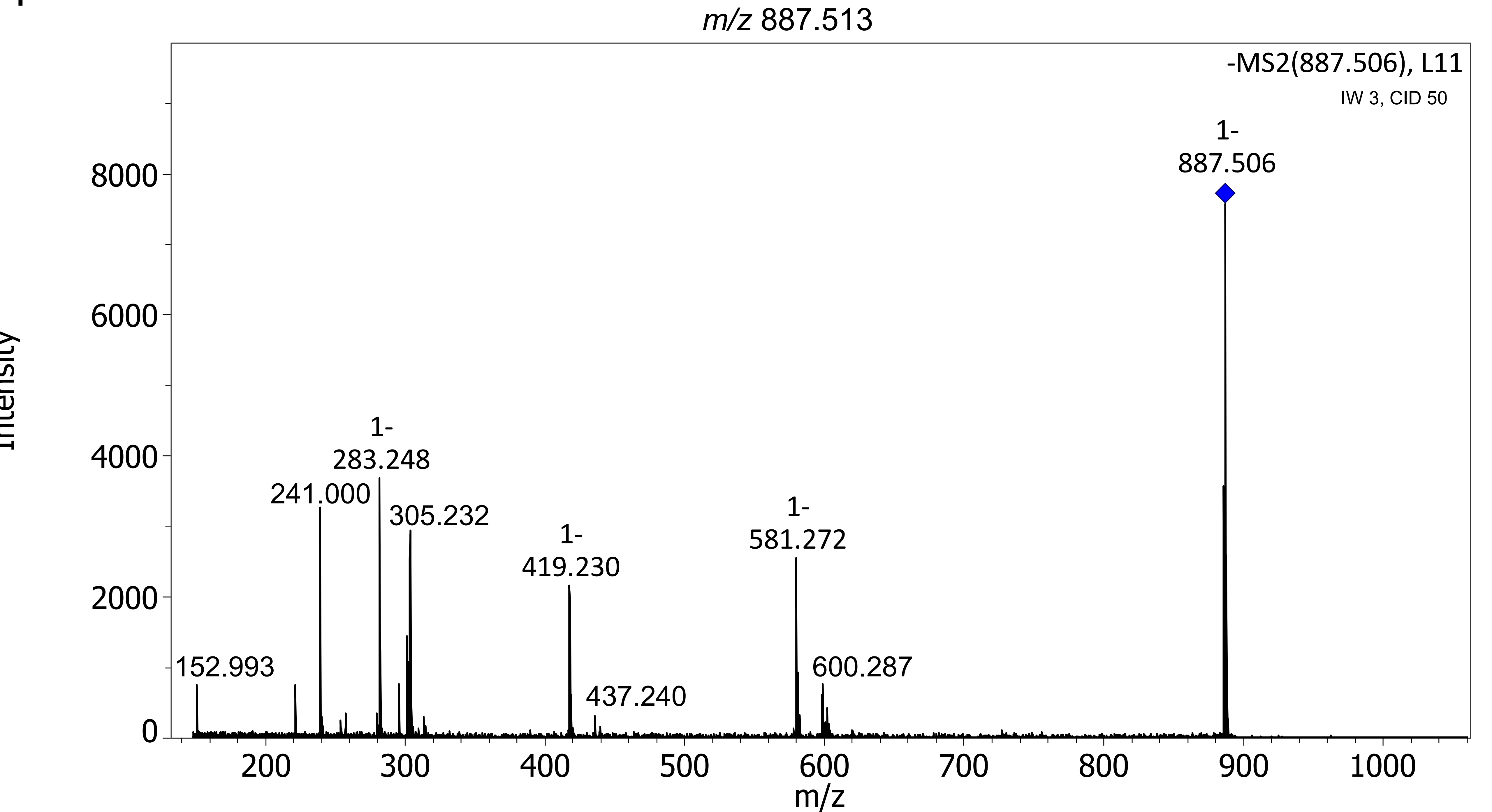

m/z 887.513
-MS2(887.506), L11
IW 3, CID 50
1-
887.506
1-
283.248
241.000
305.232
1-
419.230
1-
581.272
152.993
437.240
600.287
Intensity
m/z
0
2000
4000
6000
8000
200
300
400
500
600
700
800
900
1000

**z**

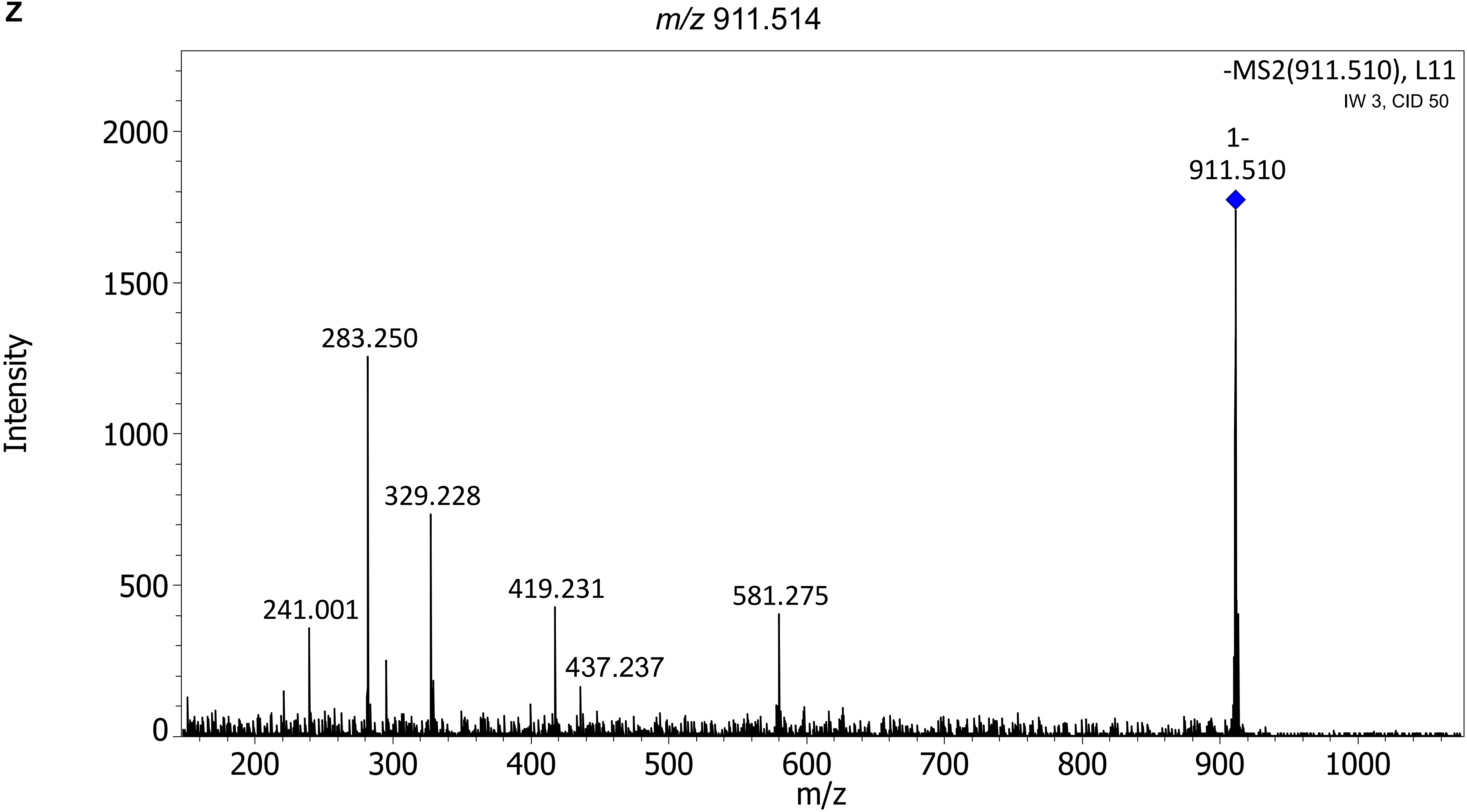

m/z 911.514
-MS2(911.510), L11
IW 3, CID 50
1-
911.510
283.250
329.228
241.001
419.231
437.237
581.275
2000
1500
1000
500
0
Intensity
200
300
400
500
600
700
800
900
1000
m/z
